# MaxHiC: robust estimation of chromatin interaction frequency in Hi-C and capture Hi-C experiments

**DOI:** 10.1101/2020.04.23.056226

**Authors:** Hamid Alinejad-Rokny, Rassa Ghavami Modegh, Hamid R. Rabiee, Narges Rezaie, Kin Tung Tam, Alistair R. R. Forrest

## Abstract

Hi-C is a genome-wide chromosome conformation capture technology that detects interactions between pairs of genomic regions, and exploits higher order chromatin structures. Conceptually Hi-C data counts interaction frequencies between every position in the genome and every other position. Biologically functional interactions are expected to occur more frequently than random (background) interactions. To identify biologically relevant interactions, several background models that take biases such as distance, GC content and mappability into account have been proposed. Here we introduce MaxHiC, a background correction tool that deals with these complex biases and robustly identifies statistically significant interactions in both Hi-C and capture Hi-C experiments. MaxHiC uses a negative binomial distribution model and a maximum likelihood technique to correct biases in both Hi-C and capture Hi-C libraries. We systematically benchmark MaxHiC against major Hi-C background correction tools and demonstrate using published Hi-C and capture Hi-C datasets that 1) Interacting regions identified by MaxHiC have significantly greater levels of overlap with known regulatory features (e.g. active chromatin histone marks, CTCF binding sites, DNase sensitivity) and also disease-associated genome-wide association SNPs than those identified by currently existing models, and 2) the pairs of interacting regions are more likely to be linked by eQTL pairs and more likely to link known regulatory features than any of the existing methods. We also demonstrate that interactions between different genomic region types have distinct distance distribution only revealed by MaxHiC. MaxHiC is publicly available as a python package for the analysis of Hi-C and capture Hi-C data.

**Author summary:** MaxHiC is a robust machine learning based tool for identifying significant interacting regions from both Hi-C and capture Hi-C data. All the current existing models are designed for either Hi-C or capture Hi-C data, however we developed MaxHiC to be applicable for both Hi-C and capture Hi-C libraries (two different models have been used for Hi-C and capture Hi-C data). MaxHiC is also able to analyse very deep Hi-C libraries (e.g., MicroC) without any computational issues. MaxHiC significantly outperforms current existing tools in terms of enrichment of interactions between known regulatory regions as well as biologically relevant interactions.

## Background

Hi-C is a genome-wide chromosome conformation capture technology used to identify interactions between pairs of genomic regions. Hi-C libraries consist of a pair of reads which can be mapped to the genome for identifying pairs of interacting regions. The data is typically summarized as a contact matrix, in which each row and column correspond to a (fixed-bin) fragment of the genome and each cell shows the number of reads supporting an interaction between the two fragments. Hi-C data has multiple biases due both to the multiple steps used in the protocol and the proclivity of closely located genomic regions that are more likely to randomly interact. Not all interactions are biologically relevant and thus the important challenge in the analysis of Hi-C data is to distinguish between random/artefactual interactions and those interactions that are more likely to be functional/regulatory.

There are two main approaches used to identify significant interactions in Hi-C data. The first, as used in Hi-C-Norm (1) and Hi-C-DC (2), attempts to quantify known sources of biases (fragment length, GC content and mappability), and use them as variables in a background expectation function. The second approach, as used in ICE (3), Fit-Hi-C (4), CHiCAGO (5) and GOTHiC (6), assumes that all Hi-C biases are reflected in the read count data of interactions. Therefore, the observed read counts are used to assign a P-value to interactions. In order to account for unknown sources of bias a separate bias parameter is assumed for each bin. The tools using the first approach are limited to analysis of Hi-C libraries generated with restriction enzymes, whereas those using the second approach can also work with DNase Hi-C experiments in which the fragment sizes are unknown. Rao *et al.* (7) also proposed HiCCUPS, which is not a background correction model, but can be used to identify chromatin loops. HiCCUPS identifies clusters of Hi-C contact matrixes and reports the centroid of the cluster, in which the frequency of the contact matrix is significantly higher than the local background. Using a local background and reporting only the centroid of a cluster results in identification of a small number of significant Hi-C interactions. Notably, as HiCCUPS is implemented using GP-GPUs, the user needs to have specific hardware to run it.

We introduce MaxHiC (Maximum Likelihood estimation for Hi-C), a negative binomial model that uses a maximum likelihood technique to correct the complex combination of known and unknown biases in both Hi-C and capture Hi-C libraries. Benchmarking against current leading background models (GOTHiC, CHiCAGO, Fit-Hi-C, Fit-Hi-C2 (8) and HiCCUPS) shows that MaxHiC identifies interactions that contain substantially more evidence of biological relevance. Interacting regions identified by MaxHiC are significantly (P-value < 0.05) enriched for H3K27ac, H3K4me1, CTCF binding sites, DNase hypersensitive sites, and GWAS SNPs. They are also more likely to be annotated as promoters, enhancers, and insulator elements. More importantly the interacting pairs are substantially enriched for interactions between regulatory regions and non-coding polymorphisms linked by eQTL to target genes.

MaxHiC as an open-source python package is available https://github.com/bcb-sut/MaxHiC.

## Results

### A negative binomial regression background model for Hi-C data

Here we present MaxHiC, a novel background correction model for identifying statistically significant interacting regions in both general and capture Hi-C libraries. In MaxHiC, the genome is divided into fixed size non-overlapping bins, and the read counts supporting interactions between bins are tested for significance. The read-count of interactions are assumed to follow a negative binomial distribution which is widely used to model over dispersed count data (9, 10). The mean parameter of the negative binomial distribution for each interaction is calculated as a function of the bias factors of its two ends and their genomic distance.

It is known that the contact frequency between pairs of regions linked by Hi-C reads decays as a function of distance along the chromosome (11). A large fraction of Hi-C reads represent random ligations products of nearby genomic regions. In MaxHiC, distance is modelled by a function that decreases at increasing genomic distances to reach a small but constant non-zero value to account for random ligations. For *trans*-interactions, we use the same constant value as observed for distant *cis* interactions. Bias factors of bins are calculated as a function of their total read-count and mappability of their surrounding bins. All of the parameters of the model are learned by maximizing the logarithm of likelihood of the observed interactions using the ADAM algorithm (12). The model is trained in multiple iterations; in each iteration, putative interactions with the best P-values are identified and set aside. These significant interactions are ignored in the next phase of training so the trained background model won’t become biased. The same approach is also used in MaxHiC to analyze Capture-Hi-C data. The only difference is that separate sets of parameters are learned for modelling target-target, target-non-target and non-target-non-target interactions to account for the fact that interactions involving one or two of the targeted regions will be enriched to different levels compared to non-targeted regions. A schematic of the MaxHiC model for analysis of general Hi-C is shown in **Figure 1a** and a detailed explanation of both models is provided in the methods section.

**Figure 1.**
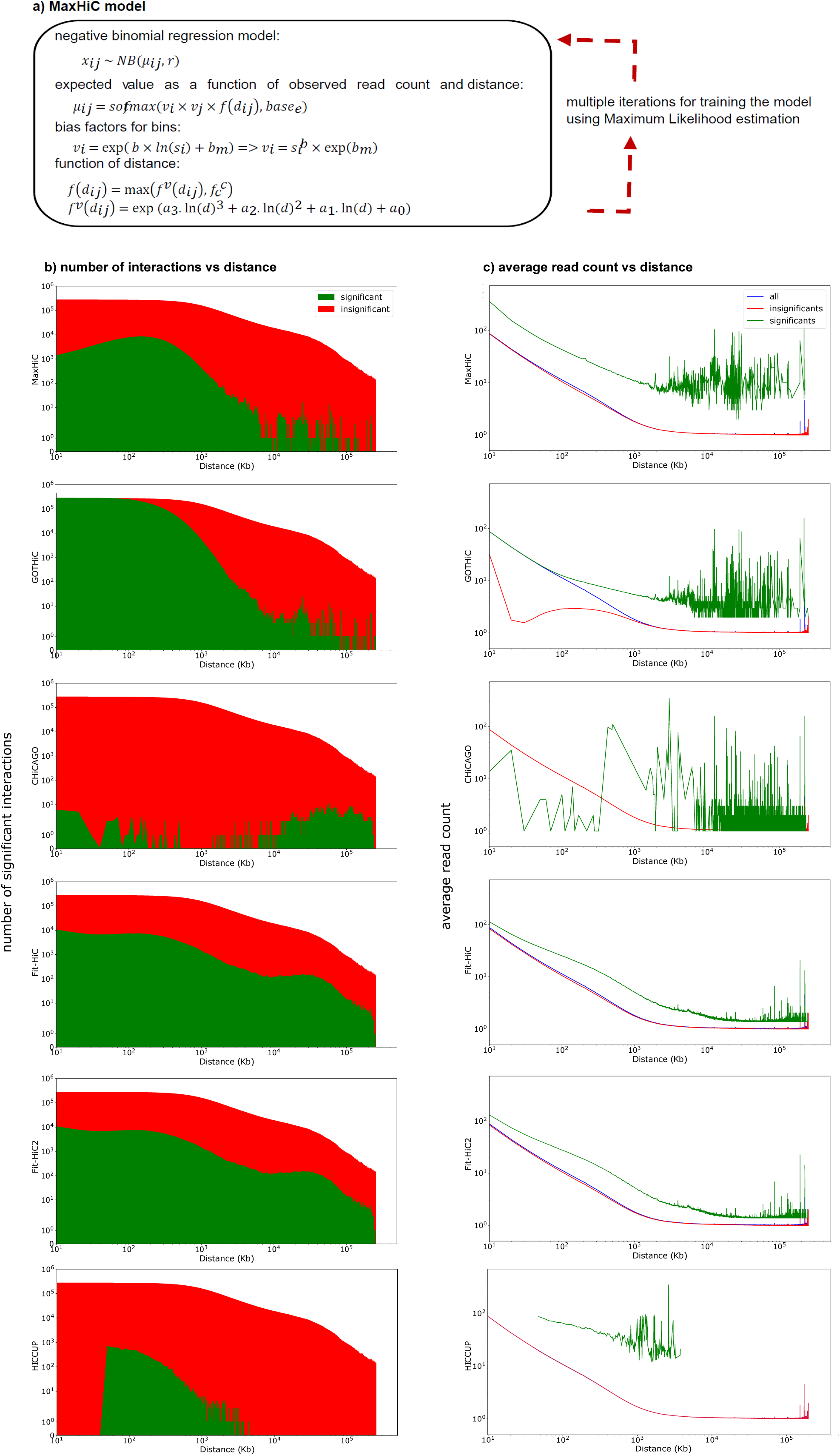
Background modelling as a function of distance in MaxHiC and other HiC tools. **a)** MaxHiC mathematical model. **b)** Average read-count of significant (green), insignificant (red), and all interactions (blue) in different genomic distances identified by different HiC tools at 10kb bin size on the Rao *et al.* GM12878 data. **c)** Number of significant interactions identified by the six models in different genomic distances on Rao *et al.* GM12878 sample at fragment size 10kb. HICCUPS doesn’t have any significant interactions in in less than 50kb or more than 5Mb. Note, we used a modified version of CHiCAGO to be applicable for non-capture Hi-C data.

### MaxHiC vs other leading Hi-C tools

In order to evaluate the performance of MaxHiC, we performed a comprehensive benchmarking between MaxHiC, GOTHiC, CHiCAGO (for all non-capture Hi-C data, we used a modified version of CHiCAGO to be applicable for non-capture Hi-C data), Fit-Hi-C, Fit-Hi-C2 and HiCCUPS (7) on several published Hi-C datasets (see more details about the samples in **Supplementary table S1**).

We first used MaxHiC to identify significant interactions (at 1kb, 5kb and 10kb bin sizes) in two publicly available Hi-C data from Rao *et al.* (7) (on the GM12878 EBV transformed B-lymphoblastoid cell line and human mammary epithelial cells (HMEC)) and one set capture Hi-C data from Mifsud *et al*. (13) (also on GM12878). For each library we also used GOTHiC, CHiCAGO, Fit-Hi-C, Fit-Hi-C2, and HiCCUPS with their default parameters to identify significantly interacting regions. **Table 1** and **Supplementary table S2** summarise the numbers of significantly interacting pairs found using each method. **Supplementary figure S1** shows Venn diagrams of the interacting pairs identified by each method at 10kb bin sizes. Notably the number of interactions called as significant by each method varied greatly (from 10,485 using HiCCUPS, to 8,136,100 using GOTHiC).

**Table 1.** Statistical summary of significant interactions identified by MaxHiC, GOTHiC, CHiCAGO, Fit-Hi-C, Fit-HiC2, and HiCCUPS in the GM12878 Hi-C library from Rao *et al.* (7). at bin-sizes 5kb and 10kb. Statistical summaries for HMEC library are provided in **Supplementary tables S2**, respectively.

| 1kb | MaxHiC | GOTHiC | CHiCAGO | Fit-Hi-C | Fit-Hi-C2 | HiCCUPS* | All |
| --- | --- | --- | --- | --- | --- | --- | --- |
| percentage of significant interactions | 0.03 | 9.4 | - | 16.6 | 17.1 | - | 100 |
| average read-count of interactions | 18.1 | 2.9 | - | 1.1 | 1.5 | - | 1.2 |
| min read-count of cis interactions | 4 | 2 | - | 1 | 1 | - | 1 |
| median distance of cis interactions (kb) | 3 | 14 | - | 30832 | 29426 | - | 544 |
| **5kb** |  |  |  |  |  |  |  |
| percentage of significant interactions | 0.17 | 9.25 | 0.17 | 0.8 | 0.9 | 9.40E-05 | 100 |
| average read-count of interactions | 21.3 | 7.85 | 1.29 | 5.6 | 5.9 | 32.39 | 1.77 |
| min read-count of cis interactions | 3 | 2 | 1 | 1 | 1 | 11 | 1 |
| median distance of cis interactions (kb) | 130 | 85 | 101350 | 5235 | 5014 | 110 | 15173 |
| **10kb** |  |  |  |  |  |  |  |
| percentage of significant interactions | 0.32 | 7.32 | 0.02 | 1.06 | 1.1 | 9.25e-05 | 100 |
| average read-count of interactions | 29.4 | 15.17 | 1.83 | 12.3 | 13.5 | 58.62 | 2.38 |
| min read-count of interactions | 3 | 2 | 1 | 2 | 2 | 11 | 1 |
| median distance of cis interactions (kb) | 270 | 170 | 93213 | 7590 | 7426 | 150 | 6990 |

\* HiCCUPS calls interactions with FDR < 0.05 as significant. For all other methods we used a threshold of P-value < 0.001 to identify significant interactions.

The sets of interactions called as significant or insignificant by each method (**Figure 1b** for sample GM12878 and **Supplementary figure S2a** for HMEC) had distinct distance distributions. At 10kb bin size, GOTHiC and HiCCUPS had the smallest median distances between interacting regions of 170kb, while that for MaxHiC, Fit-Hi-C, Fit-Hi-C2 and CHiCAGO were 270kb, 7,590kb, 7,426kb and 93,213kb respectively. The low median length observed for GOTHiC is reflected in the observation that it calls almost all putative interactions below 100kb as significant.

Comparing the interactions called as significant by each method revealed interactions identified by MaxHiC had substantially more read support than those identified by all methods except HiCCUPS, however HiCCUPS only predicted significant interactions for a small range of distances ∼50kb to ∼5000kb. Both the average number of reads and minimum number of reads supporting a significant interaction were higher for those called by MaxHiC than other models (**Table 1, Supplementary tables S2**). By examining the average read count of significant and insignificant interactions called by each method (**Figure 1c** for GM12878 and **Supplementary figure S2b** for HMEC), it is revealed that at all distances, significant interactions called by MaxHiC and Fit-Hi-C had substantially more reads than those called as insignificant. GOTHiC only demonstrated such trend at distances above 100kb. At all distances, paired regions called as significantly interacting by MaxHiC had higher average read count support than the other five methods. This was also reflected in the minimum number and average number of reads required to call an interaction as significant (**Supplementary figure S3**). We observed similar pattern when considering interactions calculated using different fragment sizes (**Supplementary figure S4** shows the same plots using 1kb, 5kb, 10kb, and 50kb bin sizes). Note: HiCCUPS was not able to run on 1kb bin size and found no loops when run on 50kb bin size.

### Evidence of regulatory potential for interacting regions identified by MaxHiC

We next investigated whether interacting regions identified by each method as significant were enriched for genomic and epigenetic annotations indicative of regulatory regions. Comparison of the interacting regions with histone mark profiles generated for GM12878 and HMEC by the ENCODE consortium (14–16) revealed the regions identified by MaxHiC were significantly (P-value < 0.05) enriched for H3K9ac, H3K27ac, H3K79me2, H3K4me3, H3k4me1, and H4K20me1 marks (**Figure 2**). They were also enriched for DNAse hypersensitive sites and CTCF binding but not for the repressive mark H3K27me3. Importantly there was very little if any enrichment of these features in regions called as significant by GOTHiC, CHiCAGO, Fit-Hi-C, and Fit-Hi-C2. Interestingly HiCCUPS had similar levels of enrichment however MaxHiC identified 34 fold more interactions. For the GM12878 analysis we also observed significant enrichment (P-value < 0.05) of transcription factor binding sites from ENCODE ChIP-seq data (**Supplementary figure S5 and Supplementary table S3**). Again, MaxHiC and HiCCUPS showed similar levels of enrichment for GM12878 specific transcription factor binding sites. Together this suggests that both MaxHiC and HiCCUPS identify interactions between regions with regulatory potential however MaxHiC identifies substantially more (34 fold).

**Figure 2.**
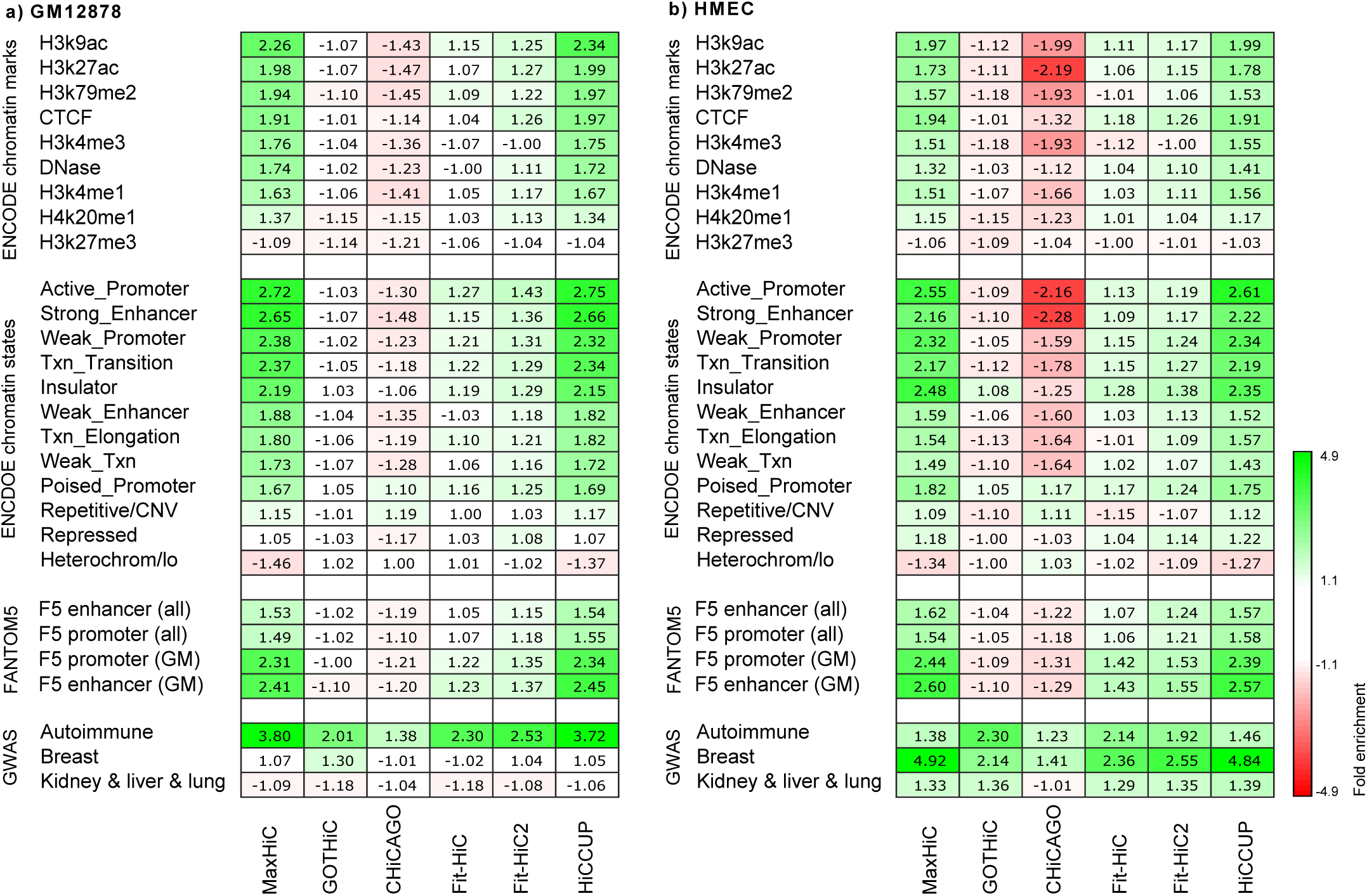
Enrichment of regulatory potential in regions identified by MaxHiC and other tools. **a)** Enrichment of GM12878 specific histone marks, putative regulatory regions and GWAS signal in regions identified by MaxHiC in the Rao *et al.* GM12878 dataset. **b)** As in (a), but enrichment of signals in the HMEC dataset is shown. ENCODE chromatin states are from GM12878 and HMEC data respectively. For FANTOM5, enrichment is shown for all FANTOM5 promoters and enhancers and for those only active in GM12878 and HMEC. GWAS SNPs for autoimmune disease, neurological/behavioural traits and kidney/liver/lung disorders (**Supplementary table 4**) were downloaded from Maurano *et al*. (26). Note: HiCCUPS calls interactions with FDR < 0.05 as significant. For all other methods we used a threshold of P-value < 0.001 to identify significant interactions.

Next, the interacting regions identified by MaxHiC were significantly enriched (P-value < 0.05) for ENCODE annotated enhancers, promoters, insulators and transcription states (17, 18) (**Figure 2**). They were also significantly depleted (P-value < 0.05) of regions annotated as heterochomatin. Again, HiCCUPS showed similar levels of enrichment while Fit-Hi-C and Fit-Hi-C2 showed some lower level of enrichment for some of the chromatin states. Surprisingly regions identified by CHiCAGO were depleted of these states.

Besides the regulatory regions predicted by ENCODE, significant interactions identified by MaxHiC also contained a much higher fraction of promoters and enhancers identified by FANTOM5 as active in GM12878 and HMEC (in comparison to all FANTOM5 promoters and enhancers) (**Figure 2**) indicating that MaxHiC not only enriches for promoters and enhancers but those active in the cell type from which the Hi-C data was generated.

Importantly, interacting regions identified by MaxHiC and HiCCUPS on the GM12878 dataset were significantly (P-value < 0.05) more likely to overlap autoimmune SNPs identified in GWAS while those identified in the HMEC were significantly (P-value < 0.05) more likely to overlap breast trait associated SNPs (**Figure 2,** GWAS traits considered are provided in **Supplementary table S4**). No such enrichment was observed for SNPs associated with unrelated traits such as kidney, lung, and liver traits.

Lastly, we compared interactions found by MaxHiC to those from the top pixel based method Fit-HiC2 and the loop caller HICCUPS. In the comparison to Fit-Hi-C2, the interactions found by MaxHiC alone were significantly enriched (P-value < 0.05) for interactions between regulatory regions (and depleted of heterochromatin and H3K27me3) whereas those found by Fit-Hi-C2 alone showed no enrichment and were near the background expectation of 1 (**Supplementary Figure S6a**). In contrast the MaxHiC and HICCUPS -specific interactions were enriched to similar levels (**Supplementary Figure S6b**). We also provided an example of Hi-C interaction identified by MaxHiC as significant, which was not identified as a significant interaction by other tools. Both sides if the interaction have strong signals of regulatory features. (**Supplementary Figure S7**).

### MaxHiC enriches for interactions between regulatory regions

We next examined the fractions of interacting pairs identified by each method involving regulatory elements at both ends. We specifically focused on promoter-enhancer, promoter-promoter, enhancer-enhancer and insulator-insulator pairs. In the GM12878 dataset approximately 12% of the raw unfiltered pairs fell into at least one of the above categories (**Figure 3a**). Strikingly, the read pairs identified by MaxHiC were almost 4 times (47%) more likely to fall into one of these four categories (**Figure 3a**). Notably ∼17% of the pairs corresponded to promoter-enhancer interactions, ∼15% enhancer-enhancer, ∼8% promoter-promoter and ∼7% to insulator-insulator interactions. Similar levels of enrichment were observed for HiCCUPS, but as noted above HiCCUPS identified 34 fold fewer significant interactions. In contrast, the other methods had distributions similar to the unfiltered pairs with Fit-Hi-C, Fit-Hi-C2 and GOTHiC showing fractions slightly above that of the unfiltered pairs. Extending this to any interaction involving an enhancer, promoter or insulator increased the fraction of annotated pairs to 70%. Similar fractions were observed with the HMEC data. **Figure 3b** shows the relative enrichment for interactions involving pairs of chromatin features and predicted chromatin states. Similar to **Figure 2** we observed enrichment of interactions involving active chromatin states and depletion of heterochromatin-heterochromatin pairs using the interactions called as significant by MaxHiC and HiCCUPS and to a much lesser extent with Fit-Hi-C and Fit-Hi-C2.

**Figure 3.**
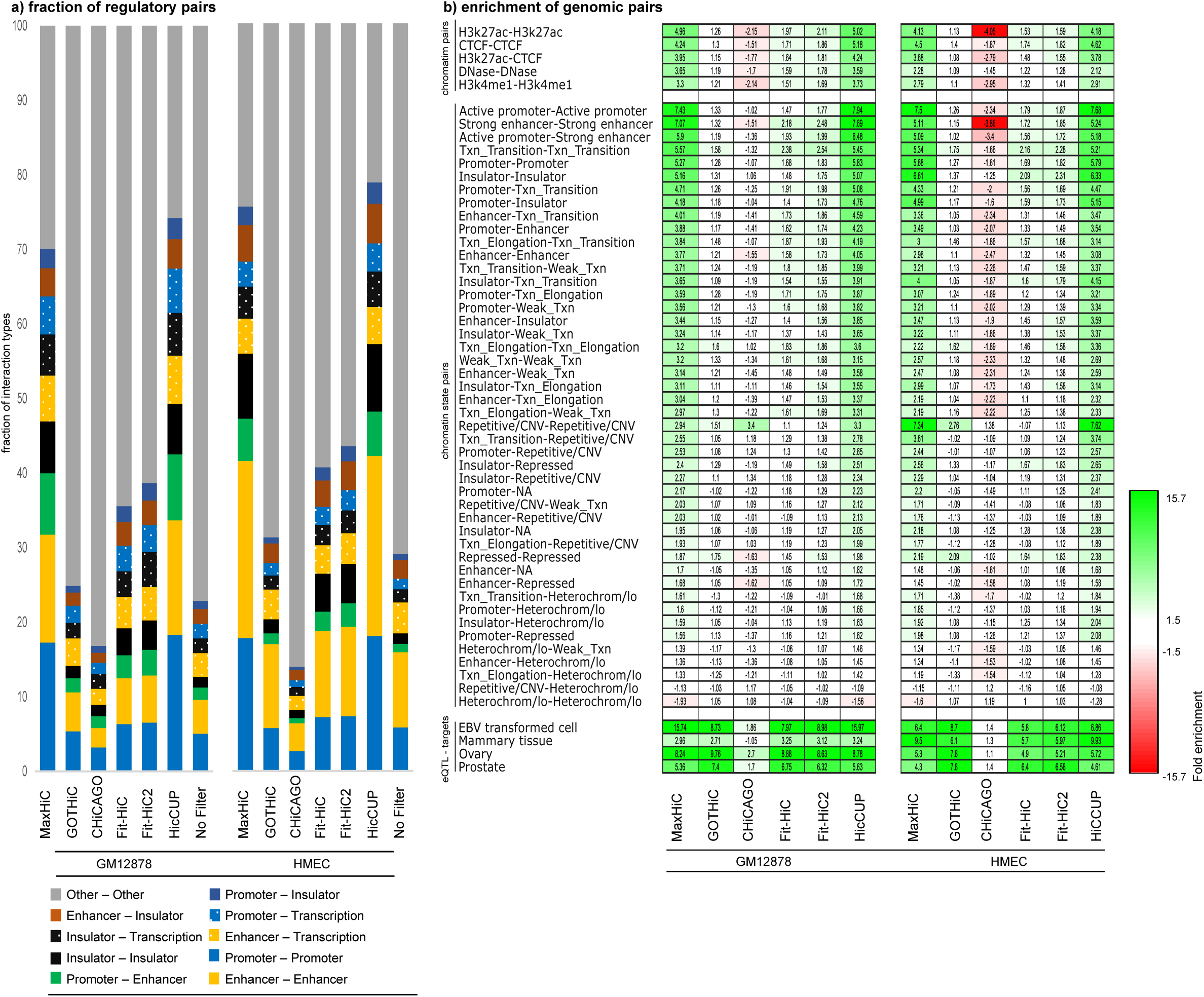
MaxHiC enriches for interactions between regulatory regions, eQTL-target pairs and known enhancer-promoter pairs. **a)** Fraction of regulatory elements in the significant interactions identified by the four models. **b)** Enrichment of cell line specific chromatin pairs, chromatin states pairs, and eQTL pairs that overlapped with both ends of the significant interactions identified by the six models at fragment size 10kb. The enrichment is calculated by dividing of the fraction of the features overlapping with both ends of the significant interactions identified by each model on fraction of the features overlapping with both ends of all interactions (*see methods*).

We next used expression quantitative trait loci (eQTL) pairs, identified in EBV-transformed lymphocytes as part of the Genotype-Tissue Expression (GTEx) Project (19) to examine enrichment of eQTL SNPs in regions that interact with promoters. Approximately 2.1% of the interactions identified by MaxHiC in the GM12878 cell line overlapped at least one eQTL pair. This corresponded to a 15.7 fold enrichment (**Figure 3b**). In contrast GOTHiC, CHiCAGO and Fit-Hi-C’s interactions were lower (8.3, 1.8 and 7.9 fold enriched respectively). Matching analyses on HMEC confirmed similar enrichments for regulatory features and eQTLs from mammary tissue (**Figure 3b**).

### Genomic spacing of interacting regulatory elements

Using the putative regulatory region annotations from the analysis above, we next examined the spacing preferences for interactions between different regulatory features. We first examined preferred spacing for H3K27ac-H3K27ac and CTCF-CTCF structural loops. For MaxHiC, Fit-Hi-C, and Fit-Hi-C2 we observed the spacing between CTCF pairs to be larger (130kb) than H3K27ac-H3K27ac pairs (80kb) (**Figure 4a**). In both cases there were long tails reaching beyond 1Mb. Performing the same analysis on annotated regulatory regions revealed promoter-promoter, enhancer-promoter, and enhancer-enhancer pairs had almost identical spacing preferences (with median spacings of 70kb, 70kb and 90kb respectively), while insulator-insulator pairs were considerably further apart with a median spacing of 170kb (**Figure 4b**). We did not see this trend for HiCCUPS however we note HiCCUPS did not identify significant interactions below 50kb and above 5Mb (**Figure 1b, c**).

**Figure 4.**
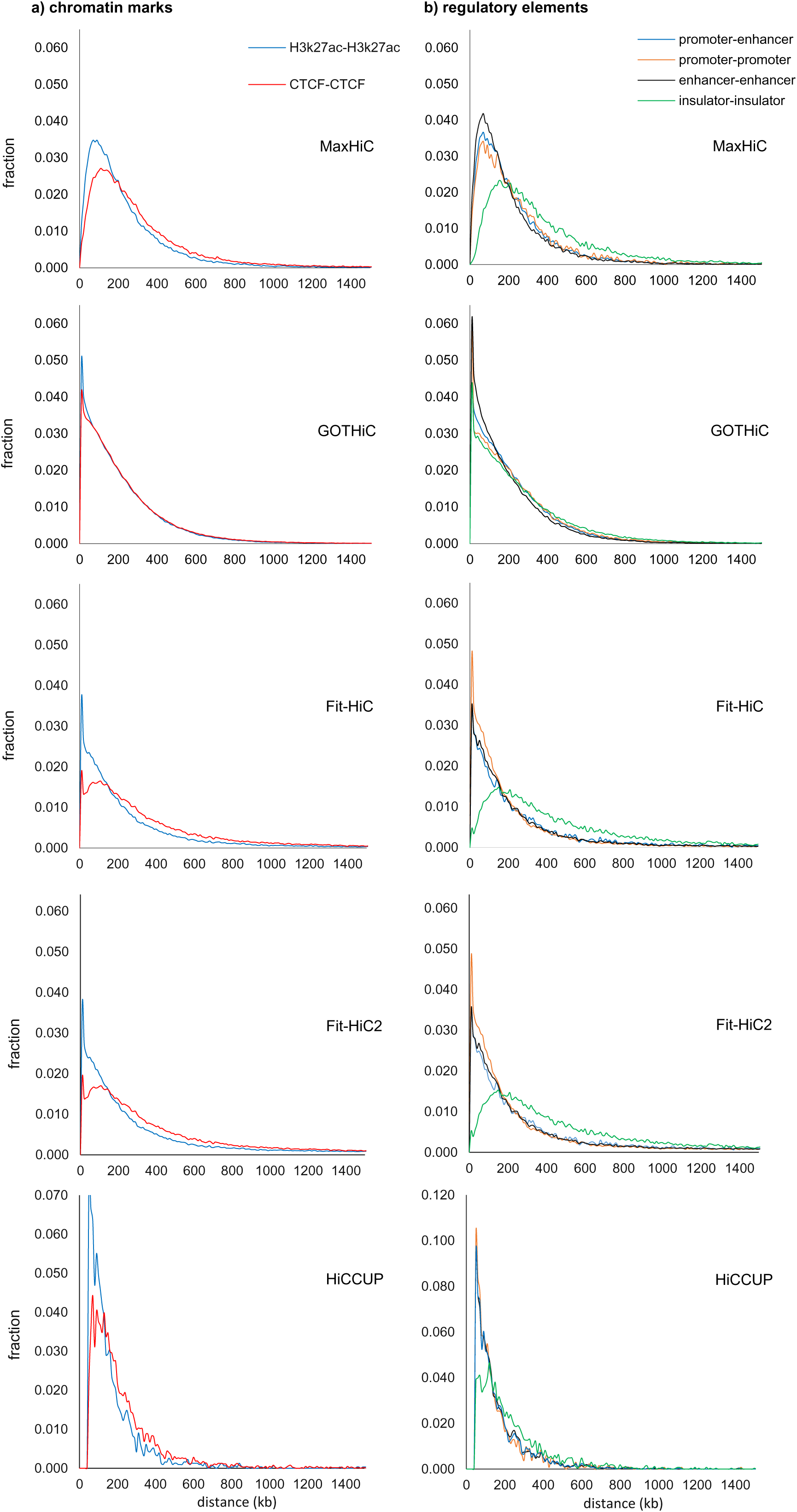
Distance distribution for interactions between different classes of regulatory elements. **a)** Comparison of spacing observed for H3k27ac-H3k27ac and CTCF-CTCF structural loops in the interactions identified by MaxHiC in the Rao *et al.* GM12878 dataset. **b)** Comparison of spacing observed for promoter-promoter, enhancer-promoter, enhancer-enhancer and insulator-insulator pairs in the interactions identified by MaxHiC in the Rao *et al.* GM12878 dataset. Plots use fragment bin-size 5kb and window size 5kb.

### MaxHiC contact maps and aggregate peak analyses

Comparing contact maps of significant interactions found for MaxHiC and HICCUPS models (**Supplementary Figure S8**) shows at first glance they are very similar, however in a zoomed in region we can see additional interactions called by MaxHiC that are missed by HICCUPS. Examining this in more detail revealed approximately 8 fold more bases are identified as significantly interacting by MaxHiC than HICCUPS (**Supplementary Figure S9**). However, to further show that these interactions are not random we provide an aggregate peak analysis (APA) comparing signals surrounding significant peaks called by MaxHiC and HiCCUPS. **Figure 5** shows the aggregate background interaction frequency of 5 bins upstream and downstream of the significantly interacting bins. APA plots are shown for the top 5k, and 10k. As the figure shows, MaxHiC and HiCCUPS have almost the same APA patterns and confirmed that the significant interactions identified by MaxHiC are not random and that even when all 352k significant interactions identified by MaxHiC are considered there is strong focal enrichment in the APA. To demonstrate MaxHiC application to a large dataset we also analysed a MicroC dataset (20) consisting of ∼480M raw valid interactions. The contact map for this dataset is shown in **Supplementary Figure S10a**. Reassuringly matching aggregate peak analysis plots confirm there is strong focal enrichment for these significant interactions **Supplementary Figure S10b**.

**Figure 5.**
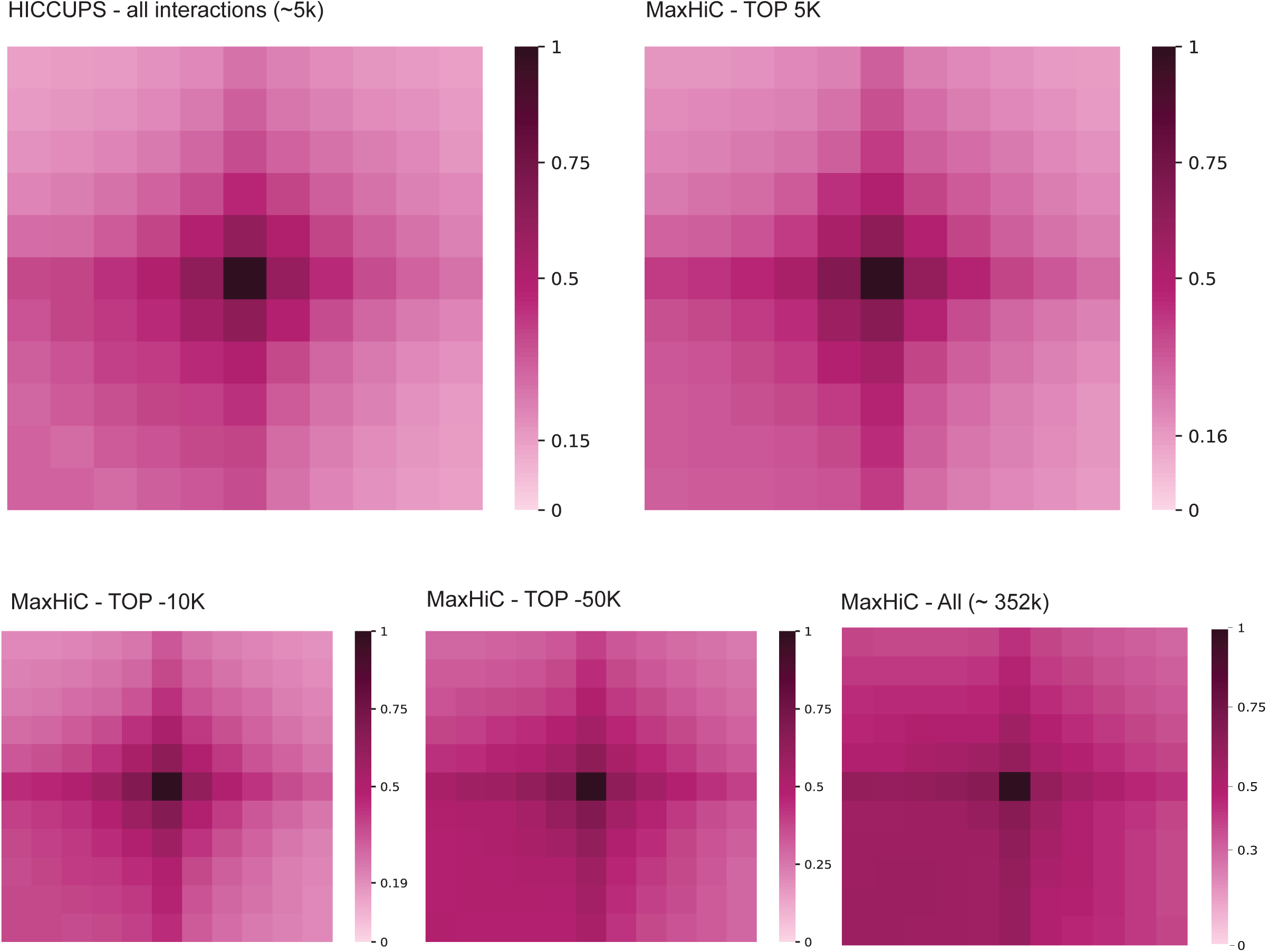
Aggregate peak analysis (APA) of significant interactions called by MaxHiC and HICCUPS. Interactions were called at 10kb resolution using the GM12878 Hi-C dataset from Rao *et al*. The analysis shows the aggregate background interaction frequency of 5 bins upstream and downstream of the significantly interacting bins. For HICCUPS the bins at the centroid of the loop regions are used. For HICCUPS the APA for all 5,348 interactions are shown. For MaxHiC, APA plots are shown for the top 5k, 10k, 50k and all 243k significant interactions.

### Application of MaxHiC to identify differentially interacting regions

To further demonstrate the use of MaxHiC in comparative Hi-C experiments we applied it to Hi-C datasets from cells depleted of the cohesion release factor WAPL (Haarhuis *et al*. (21)) and the polycomb protein RING1B cells (Boyle *et al*. (22)); and their matched wildtype controls. In the case of WAPL-depleted HAP1 cells MaxHiC identified almost 3 times more significant interactions than in matched wildtype cells (**Figure 6a,b**). Furthermore the distance distribution of interacting genomic regions found in the ΔWAPL cells (median = 6.4Mb) was significantly longer in comparison to wildtype (median = 3.1Mb) (**Figure 6c**); this reiterates the finding of Haarhuis *et al*. (21).

**Figure 6.**
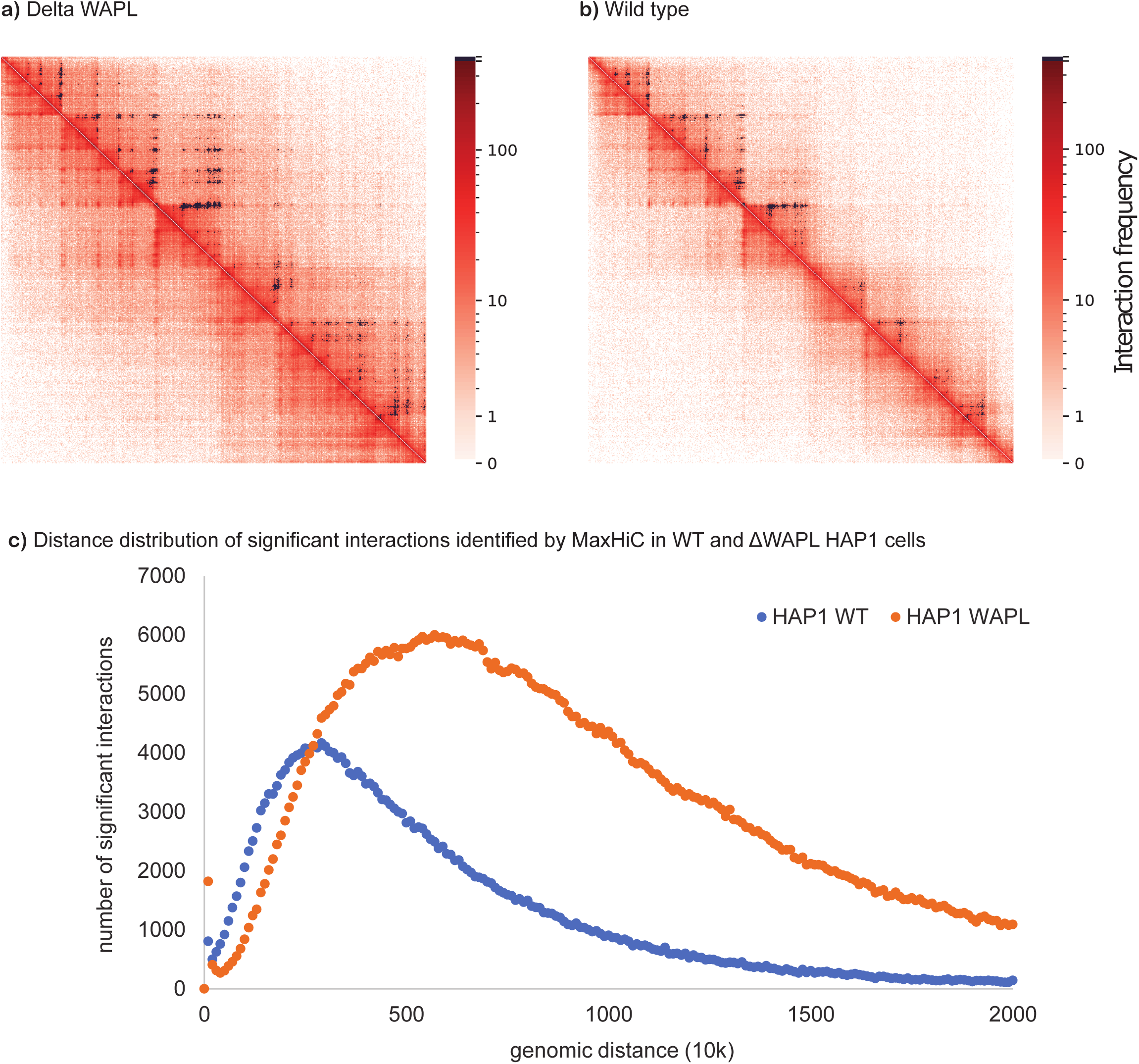
WAPL restricts chromatin loop extension. HiC Contact map comparing significant interactions identified in WT (**a**) and ΔWAPL HAP1 (**b**) cells at 10kb fragment size. Black pixels are significant interactions identified by MaxHiC at P-value <0.001. (chr7: 25M-31M). **c)** Distance distribution of significant interactions identified by MaxHiC (P-value <0.001, 10kb bins) in WT and ΔWAPL HAP1 cells. The number of significant interactions identified in HAP1 WT is 283,028 and the number of significant interactions identified in HAP1 WAPL is 761,820.

In the case of the RING1B depleted cells, MaxHiC was able to identify differentially interacting regions identified in the original paper (**Supplementary Figure S11a-S11b**; shows replicates Figure 5a from Boyle *et al*. (22)). We also observed that wildtype cells have a slightly larger median distance between interactions (325kb for WT) than the knockout cells (258kb for RING1B−/−) (**Supplementary Figure S11c**). Lastly we examined enrichment for repressive H3K27me3 and RING1B sites (using ChIP-seq data from wildtype cells (23)) in interacting regions found in wild type cells and RING1B−/− cells. In all cases the enrichment was close to the genome wide background expectation of 1, however the interactions found in the knockout were slightly more enriched than the wildtype, which suggests the sites are more open for interactions after removal of RING1B (**Supplementary Figure S11d**).

## Discussion

Enhancers have important roles in controlling gene expression. Linking distal regulatory regions such as enhancers to the genes that they regulate is critical to the interpretation of regulatory variants identified in whole genome data. Importantly, genomic variants in enhancers and regulatory elements have been linked to human complex diseases (24, 25). Therefore, pairing enhancer regions to their regulatory targets is crucial for diagnosis of genetic diseases. Compared to simplistic assignment of enhancers to their nearest promoters, computational methods have improved the prediction of enhancer-promoter pairs (26–29). However, physical interaction data is still far more convincing in validating the pairs. Hi-C data offers one approach to identify these links, however Hi-C sequencing libraries are extremely noisy because of self-ligations, random ligations artifacts and other complex sources of biases such as GC content and mappability of the sequences. Given this, accurate identification of significantly interacting regions is dramatically affected by the tool used. Here, we have presented MaxHiC, a new tool for identifying significantly interacting regions from Hi-C data (and capture Hi-C data see **Supplementary figure S12** and **Supplementary table S5**). Through a dramatically improved discrimination of biologically relevant interactions from background, we showed that MaxHiC enriches for interacting regions with substantially more evidence of regulatory potential (**Figures 1 and 2**) than those identified by existing Hi-C analysis tools (4–6, 8). This is a critical advance as enhancer-promoter interaction maps are needed to interpret the impact of non-coding variants identified in intergenic regions.

From our analyses, only MaxHiC and HiCCUPS (and to a much lesser extent Fit-Hi-C/2) enrich for interactions between regulatory regions. Specifically, interactions identified by MaxHiC were enriched for regulatory features, disease-associated genome-wide association SNPs, eQTL pairs and experimentally validated enhancer-promoter interactions. Although interacting regions identified by HiCCUPS had similar levels of enrichment it found 34 fold less significantly interacting regions. Additionally, HiCCUPS appears to have limitations on the distances at which interactions can be found and the bin sizes possible. It also required specialised GPU architecture to run. From this observation we conclude that MaxHiC is a substantially better tool than existing Hi-C analysis packages.

We also implemented a capture version of MaxHiC to analyse capture Hi-C experiments; in the capture Hi-C technology, target regions (also called baits) are sequenced with higher depth than the others, resulting in three different types of interactions target-target, target-other and other-other with different properties. Capture Hi-C baits, but not other ends, have a further source of bias associated with uneven capture efficiency, thus for capture Hi-C experiments we model each class separately. As **Supplementary figure S12** shows the capture version of MaxHiC also identifies target-interacting regions that are enriched for regulatory features and disease-associated GWAS SNPs. The levels of enrichment are much higher than the regions identified by GOTHiC, Fit-Hi-C, and Fit-Hi-C2. Same as the non-capture Hi-C analysis, MaxHiC and HiCCUPS show similar levels of enrichment in capture data, however, MaxHiC identified 36 times more interactions than HiCCUPS (36 times more interactions at 10kb fragment; 7.3 times more interactions at 5kb fragment). Surprisingly, the average read count of significant interacting regions identified by MaxHiC were much higher than HiCCUPS’s interactions (1.4 fold and 2.4 fold in 10kb and 5kb fragments, respectively). Importantly, MaxHiC’s interacting regions had higher level of enrichment than the interacting regions identified by CHiCAGO, a specific model designed for the analysis of capture Hi-C data.

In conclusion, MaxHiC is a new open source tool for identifying significant interacting regions from Hi-C and capture Hi-C data. We have demonstrated that it significantly outperforms existing tools in terms of enrichment of interactions between known regulatory regions. We will continue to develop MaxHiC related tools for identifying topologically associating domains (TADs) and for identifying interactions that are significantly different between conditions. As single-cell Hi-C data becomes more broadly available we will also adapt MaxHiC to work with these datasets. Lastly, we believe that with minor modifications the same principles used here to analyse Hi-C data may be more broadly extended to identify significant interactions from RNA-chromatin interaction datasets (30, 31).

## Online methods

### Tool availability

The MaxHiC source code and python package, a sample dataset, and instructions on how to run MaxHiC are provided at https://github.com/bcb-sut/MaxHiC. For more details about each parameter, please visit the github page. A tutorial on using the MaxHiC package is also provided in **supplementary file 1** (instructions for MaxHiC). MaxHiC is memory and speed optimized and will work with both single core and multicore machines. It can work with very large Hi-C data sets (as bam file or contact maps generated by other tools) to identify significant interactions at a broad range of resolutions (1kb-500kb).

### Publicly available Hi-C data and data access

Samples GM12878 B-lymphoblastoid cell line (SRR1658572) and Human Mammary Epithelial Cell (SRR1658680) from Rao *et al*. (7), capture Hi-C sample GM12878 B-Lymphoblastoid (ERR4360330) from Mifsud *et al*. (13) and mouse cortex Hi-C sample (SRX128219) from Dixon *et al*. (32) were used to benchmark MaxHiC against other models. All samples are publically available. More details about the samples are provided in **Supplementary table S1**.

### Mapping, filtering and Interaction calling

MaxHiC takes as input either bam files or contact map files generated by packages such as HiCUP (33) and HiC-Pro (34). For the analyses presented here, FASTQ paired-end reads were aligned to the human hg19 genome and filtered through the HiC-Pro pipeline developed in our lab (statistics are presented in **Supplementary table S1**). To remove duplicate reads, filter for valid interactions, and generate Hi-C interaction matrices (5kb and 10kb bin-sizes), we set the following parameters: MIN_FRAG_SIZE = 230; MAX_FRAG_SIZE = 1100; MIN_INSERT_SIZE = 120; and MAX_INSERT_SIZE = 990. All experimental artifacts, such as circularized reads and re-ligations, singletons have been filtered out using HiC-Pro. We also removed all self-interactions from the downstream analyses. In-house scripts have then used to statistics evaluation of Hi-C libraries (see **Supplementary table S1**).

### The MaxHiC algorithm

In the following, we explain MaxHiC algorithm and mathematics behind the model for the analysis of both Hi-C and capture Hi-C experiments.

### A negative binomial regression background model for Hi-C data

In MaxHiC, a negative binomial distribution has been used to model the read count of interactions based on the bias factors of the two ends and their genomic distance. The negative binomial distribution is the analytic form of a Poisson distribution with its mean parameter following a gamma distribution to account for over-dispersion in data. As shown in equation 1, the observed read count of interactions, *x*_*ij*_, is modelled using their expected read count, *μ*_*ij*_, and a shared dispersion factor, *r*, between all interactions.

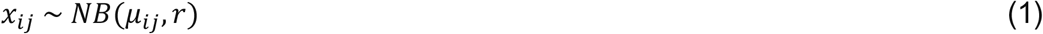

MaxHiC considers ‘neighborhooding’ (similar to local background) when identifying significant interactions. The parameters of the model are learned using a maximum likelihood approach. Gradient descent has been used to find the values of the parameters maximizing the logarithm of the likelihood function. In order to have higher accuracy, by having a higher difference between the read count of meaningful and meaningless interactions, and also to have higher speed, the proposed model uses the interactions between DNA bins, not DNA fragments. DNA is divided into pieces and all reads recorded for each fragment are assigned to the bin containing their middle points. All of the recorded interactions between pairs of bins with at least one read are the input samples used for training the model. After the model has been trained, the P-value of each interaction can be calculated based on the value of reliability, *1 - CDF*, for its observed read count.

#### Calculating the expected value for the read count

The expected read count for the interaction between bins *i* and *j* is calculated according to equation 2. *f*(*d*_*ij*_) indicates the expected value of read count to be recorded in the experiment for any two loci with distance *d*_*ij*_, *v*_*i*_ and *v*_*j*_ are bias factors of the two interacting bins. As mentioned above, bins have different characteristics and different probabilities of showing up in the experiment that affects the expected value of their interactions. It’s assumed that the effects of two bins are independent from each other, so the total bias factor has been calculated as a product of the bias factors of the two ends.

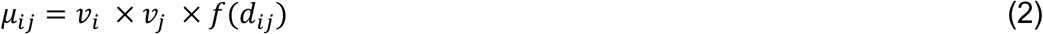

#### Calculating bias factors for bins

In order to be applicable to all types of Hi-C protocols, accounting for the unknown sources of bias and also the relation between the known sources, the second approach, mentioned in the introduction, has been adopted in this study. A uniform bias factor has been assumed for each bin, which becomes more valid as the resolution of the analysis increases and the length of bins become shorter. We expect the bins included in more interactions with higher reads to have higher tendency to show up in the experiment. In addition, the total number of interactions that a bin has been included in, would have no effect on its bias. To deal with these problems, the bias factor for bins have been calculated using equation 3 in which *v*_*base*_, *b* and *b*_*m*_ are the parameters of the function and *s*_*i*_ is the total number of reads observed for bin *i*. Here it is assumed that the total number of reads observed for each bin is a measure of its ability to show up in the final results of the test, and bias factor is assumed to be a linear function of *s*_*i*_ in log-log space. *v*_*base*_ is also considered as the base possible value for the bias factor. This prevents the bins with very few recorded reads to affect the parameters in the training process and it also prevents their related interactions with very few reads from showing up as significant because of their abnormally low bias. Many tools do this by putting a threshold and filtering bins with less bias, but in MaxHiC it is done automatically without any need for adjusting the threshold. As illustrated in equation 3, *v*_*i*_ is proportional to *s*_*i*_ raised to a power *b.* The power can be learned and if it becomes less than one it can express a saturation phenomena. As duplicated read pairs are discarded in post-processing, having more read count for two interacting bins as the read count of their interaction increases, becomes lower than their initial potential.

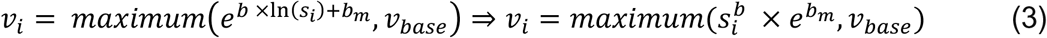

#### Calculating expected read count as a function of distance

The expected read count as a function of distance for cis interactions is illustrated in equation 4. The function is the maximum of two values. The first one, *f*^*v*^(*d*_*ij*_), is a degree 3 polynomial function of genomic distance in log-log space which specifies the expected read count of interactions based on the linear genomic distance between the two interacting loci. The reason for using an exponential function is the fact that Brownian motion and diffusion can be modelled using an exponential function. This assumption is also consistent with the power law function observed for the read count as a function of distance which has been introduced by Lieberman *et al*. for the first time (35). The degree of 3 has been used so the function can have varying curvature in different distances. In this function, the logarithm of distance has been used instead of the distance itself, as DNA is not a straight string and has multiple folding with different scales that causes proximity to have a closer relation with the logarithm of the length than the length itself. A similar concept has been used in Fit-Hi-C (4), CHiCAGO (5) and HiC-DC (2) methods. The biggest difference is the fact that all of these methods consider a strictly decreasing function for the expected read count which approaches zero as the genomic distance increases and causes all interactions between far genomic loci to be considered significant while they may have only one read. This cannot be completely logical as farther than some specific distance, random collisions occur due to three dimensional closeness not the genomic closeness. This three dimensional closeness can be due to closeness to the fixed three dimensional structures such as loops or random dynamic crossing of DNA string while two different parts of it are having contact. The expected read count for these kinds of random interactions has been modeled using a constant parameter, *f*^*c*^. Therefore, in close genomic distances *f*^*v*(*dij*)^ is dominating while the value of *f*^*v*(*dij*)^ becomes very small, the value of *f*^*c*^ dominates the expected read count. For trans interactions, similar to the cis interactions for far loci, a constant parameter has been used to denote the expected value of read count for any random trans interactions.

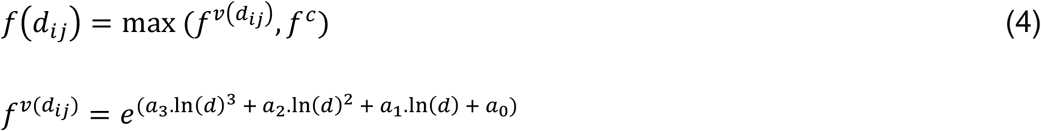

Maximum function is not a smooth function to use in optimization. Therefore, instead of maximum, the soft-maximum function has been used. This soft-maximum resembles the maximum function. When an argument, for example *α*, is larger than the other, it’s exponential becomes far more larger and so it’s related factor, 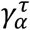, approaches 1 while the other factor, 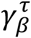, approaches 0. Therefore, the final value will be approximately equal to *α* itself.

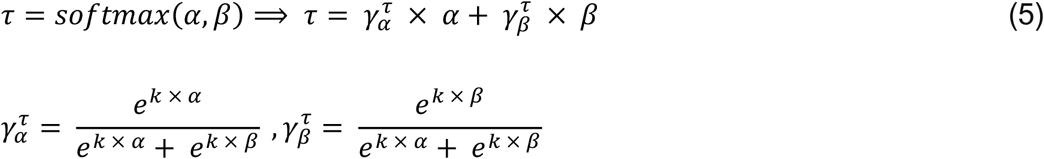

#### Training the model in multiple rounds

As mentioned above, the method models read count of interactions occurring based on Brownian motion, not the meaningful chromatin contacts, which are called real interactions from now on. But real interactions are also among the data and can affect the parameters of the model and pull them up. For example, in distances where there are more read interactions, the expectation may become higher than other distances. To deal with this bias, the model is trained in multiple rounds, which can be specified by user (the default value for iterations is 4). In each round, first the parameters of the model are learned using a stochastic gradient descent approach. The training is done in iterations until all the parameters of the model have changed by less than 0.0001 compared to the previous iteration. Training also will stop when reaching 1000 iterations to handle oscillating situations around the optimum values. This number is high enough to give the model enough time to be trained well and has been chosen by the developers based on running the model on multiple Hi-C datasets (25 samples with 2-4 different resolutions). Next, a P-value is calculated for each interaction using the trained model and the real interactions are identified based on a fixed user-defined P-value as significance limit. The read count of these real interactions is substituted with their expected read count for calculating the total number of reads for each bin, as if they were not read what would their read count be, to remove the bias of having real interactions from bias factors of the bins. They are also completely ignored in the training of the model’s parameters. This is another main difference between MaxHiC and other models.

#### An automatic way of removing the effect of noisy data

Typically, a high percentage of interacting fragments in Hi-C have only 1 read between; for example, the typical percentage can be above 85% for a resolution of 5Kb. End bins of many of these interactions have very low coverage which results in a very small expected value for the interaction. This can pull the value of parameters down for the benefit of themselves in the model. In order to cope with such noisy data, the expectation for read count is calculated based on equation 6 instead of equation 2. In this way, the maximum of expected value and a base expected read count, *base*_*e*_, is used as the final expected value. This will prevent the mentioned noisy interactions to affect greatly the parameters of the model.

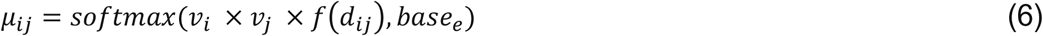

### A negative binomial regression background model for capture Hi-C data

Capture-Hi-C is a version of Hi-C in which some regions in DNA, which are called *Baits* or *Target Regions*, are sequenced with higher depth than the others. This will result in three different types of interactions in Capture-Hi-C with different properties, Target-Target, Target-Other and Other-Other. The background model developed for Capture-Hi-C follows the same statistical model as the one developed for General-Hi-C, but in order to account for the differences between these types of interactions, three different sets of parameters are used, each for one interaction type. Each set of parameters is learned separately from the others based on its own interactions, in the process of learning the parameters of the model. In calculating the bias factors for bins, the sum of read counts cannot be used directly as different interactions correspond to different types. To remove this barrier, all the read counts of interactions are converted to their equivalent read-counts in Target-Target interaction type:

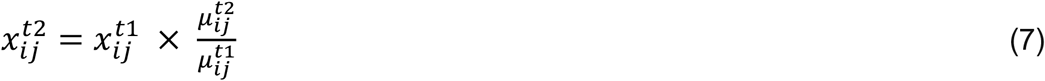

In equation 7, t1 and t2 stand for the original interaction type and the one we want to convert the read-count to, which is assumed to be Target-Target, respectively. The expected value for each interaction type is easily calculated by using the set of parameters related to that interaction type and the calculated properties of the interaction, sum of reads for bins and their genomic distance, in the expected read-count formula regardless of the original type of the interaction.

### Assessment of feature enrichment (ENCODE regulatory features, FANTOM5 promoters/enhancers)

#### Annotation datasets

We downloaded tissue-specific ChIP-Seq data in the form of processed peak calls for histone modifications (e.g. H3K27ac, H3K4me1, etc), CTCF binding sites, DNase hypersensitive sites, and ChromHMM predicted promoters and enhancers from the ENCODE project (17). We used FANTOM5 active promoters and transcribed enhancers from the FANTOM5 consortium (36, 37). FANTOM5 tissue-specific enhancers were obtained from (37).

#### Region-based analysis

To compute the enrichment of interactions overlapping an epigenetic feature, we first calculated the fraction of significant interactions with at least one end overlapping a feature. We then calculated fraction of all interactions overlapping with the feature as the genome-wide expectation. The enrichment value was then calculated by dividing the fraction of significant interactions overlapping a feature by the genome-wide expectation.

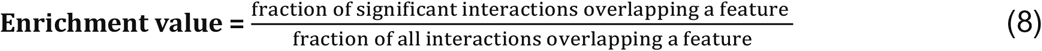

#### Pair-based analysis

To compute the enrichment of interactions that both sides of an interaction overlap with an epigenetic feature, we repeated the above analyses in which both sides of interaction overlap with the feature.

#### Capture Hi-C analysis

In the capture Hi-C analysis, to ensure our results were not affected by the strong peak signal over target regions, only target-interacting regions (“bait-to-any interactions”) are considered and signals that mapped as bait-bait interactions and those signals that mapped outside of the target-interacting regions are excluded from the analysis. To calculate the fraction of significant target-interacting regions overlapping an epigenetic feature in the capture Hi-C analysis, we first calculated the fraction of significantly target-interacting regions overlapping a feature. We then randomly selected fragments that had no interaction with the target regions (we selected exactly the same number of fragments as the number of significant target-interacting fragments). The enrichment value then calculated by dividing the fraction of significant interacting regions overlapping a feature on the average fraction of 1000 permutations. We performed the above analysis for each Hi-C background model, separately.

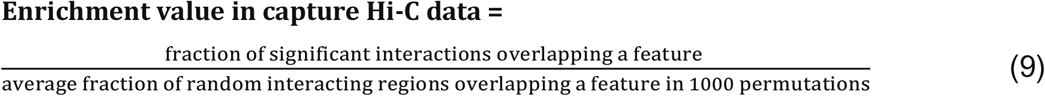

### GWAS data analysis

GWAS SNPs for autoimmune disease, neurological/behavioural traits and kidney/liver/lung disorders were downloaded from Maurano *et al*. (26). To compute the enrichment of interactions overlapping GWAS SNPs, we first calculated the fraction of significant interactions with at least one end containing GWAS SNPs. We then calculated fraction of all interactions overlapping GWAS SNPs as the genome-wide expectation. The enrichment value was then calculated by dividing the fraction of significant interactions overlapping GWAS SNPs by the genome-wide expectation. In the analysis of capture Hi-C library, we only considered overlapping of GWAS SNPs with target-interacting regions in both the significant target-interacting regions and the permutated target-interacting regions.

### Overlapping Hi-C interactions with eQTL pairs

We used the set of GTEx v7 eQTLs identified as significant in EBV-transformed lymphocytes from the Genotype-Tissue Expression (GTEx) Project (19). To calculate the fraction of eQTL pairs overlapping with significant interactions, we considered those Hi-C interactions where one side of the interaction overlapped an eQTL SNP and the other side overlapped the promoter of the eQTL target gene identified by GTEx. The promoter regions are defined as 5kb± of starting point of a gene (TSS). To calculate enrichment, we divided these by the genome-wide expectation.

### Tools running

All tools were run with their default parameters.

## Declarations

### Author contributions

HAR, ARRF, and RG designed the study. HAR, ARRF, and RG wrote the manuscript; HAR, ARRF, RG, and HRR edited the manuscript; RG implemented the tool with help from HAR. RG and HRR designed the statistical background model. HAR and RG carried out all the analyses including the statistical analyses, significant region identification, benchmarking, annotations, and text mining. HRR and NR helped with the statistical analysis. HAR and RG generated figures and tables. ARRF provided his advices in generating figures. All authors have read and approved of the final version of the paper.

### Ethics approval and consent to participate

NA

### Availability of data and materials

All data are publicly available and cited in the paper.

### Consent for publication

All authors have read and approved the final version of the paper.

### Tool availability

The MaxHiC source code and python package, a sample dataset, and instructions on how to run MaxHiC are provided at https://github.com/bcb-sut/MaxHiC.

## Acknowledgments

This work was supported by an Australian Research Council discovery project grant to ARRF (DP160101960) and a WA Department of Health Near-Miss Merit Award to HAR. ARRF is currently supported by an Australian National Health and Medical Research Council Fellowship APP1154524. ARRF was also supported by funds raised by the MACA Ride to Conquer Cancer and a Senior Cancer Research Fellowship from the Cancer Research Trust. HRR and RG were supported by IRN National Science Foundation (INSF), Grant No. 96006077. HAR is supported by a UNSW Scientia Program Fellowship. HAR was previously a research associate in Harry Perkins Institute of Medical Research under supervision of ARRF. Analysis was made possible with computational resources provided by the Telethon Kids Bioinformatics Server with funding from the Australian Government and the Government of Western Australia.

## Supplementary figure legends

**Figure S1.**
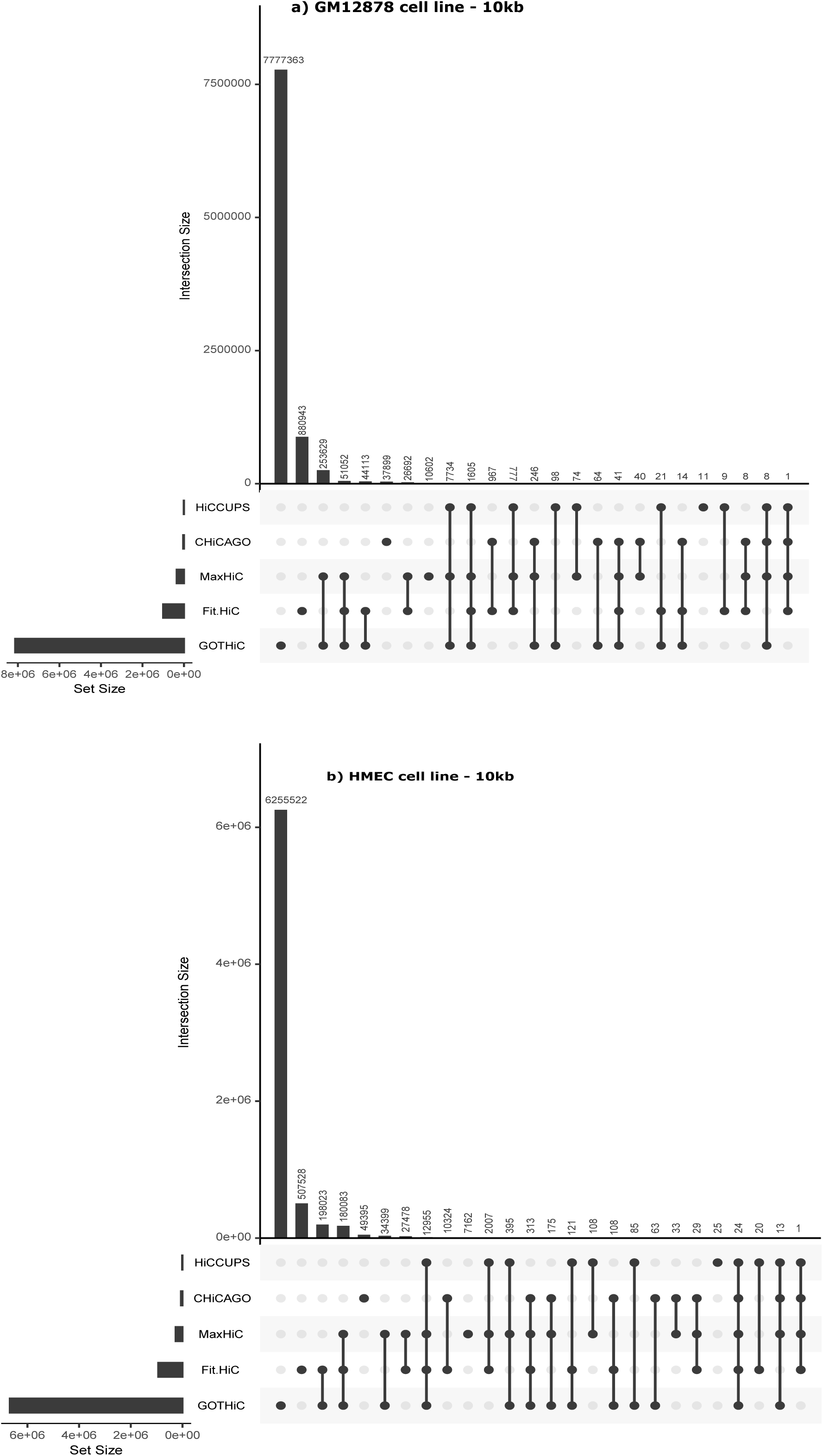
Venn diagram of the significant interactions identified by each model in the Rao *et al.* GM12878 and HMEC datasets. **a)** Venn diagram of significant interactions identified by each model in the GM12878 data at fragment size 10kb. **b)** Venn diagram of significant interactions identified by each model in the HMEC data at fragment size 10kb. We did not include Fit-Hi-C2 in this figure as it is almost the same as Fit-Hi-C.

**Figure S2.**
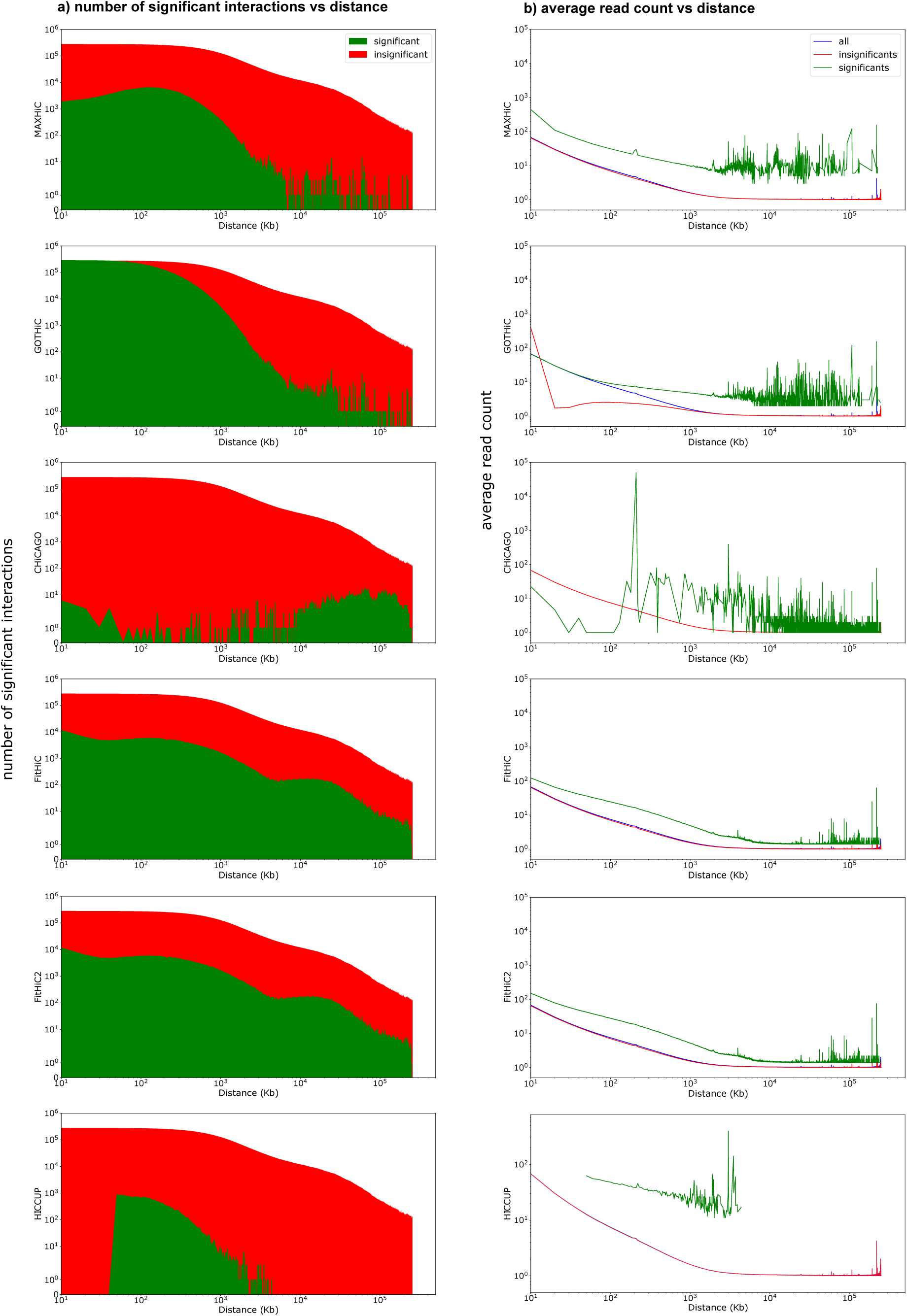
Comparisons of the number of significant interactions and average read-counts of significant interactions identified by four Hi-C background models applied to the HMEC Hi-C data. **a)** Average read-count of significant (green), insignificant (red), and all interactions (blue) in different genomic distances on the HMEC sample at fragment size 10kb. **b)** Number of significant interactions identified by the four models in different genomic distances on the HMEC sample at fragment size 10kb.

**Figure S3.**
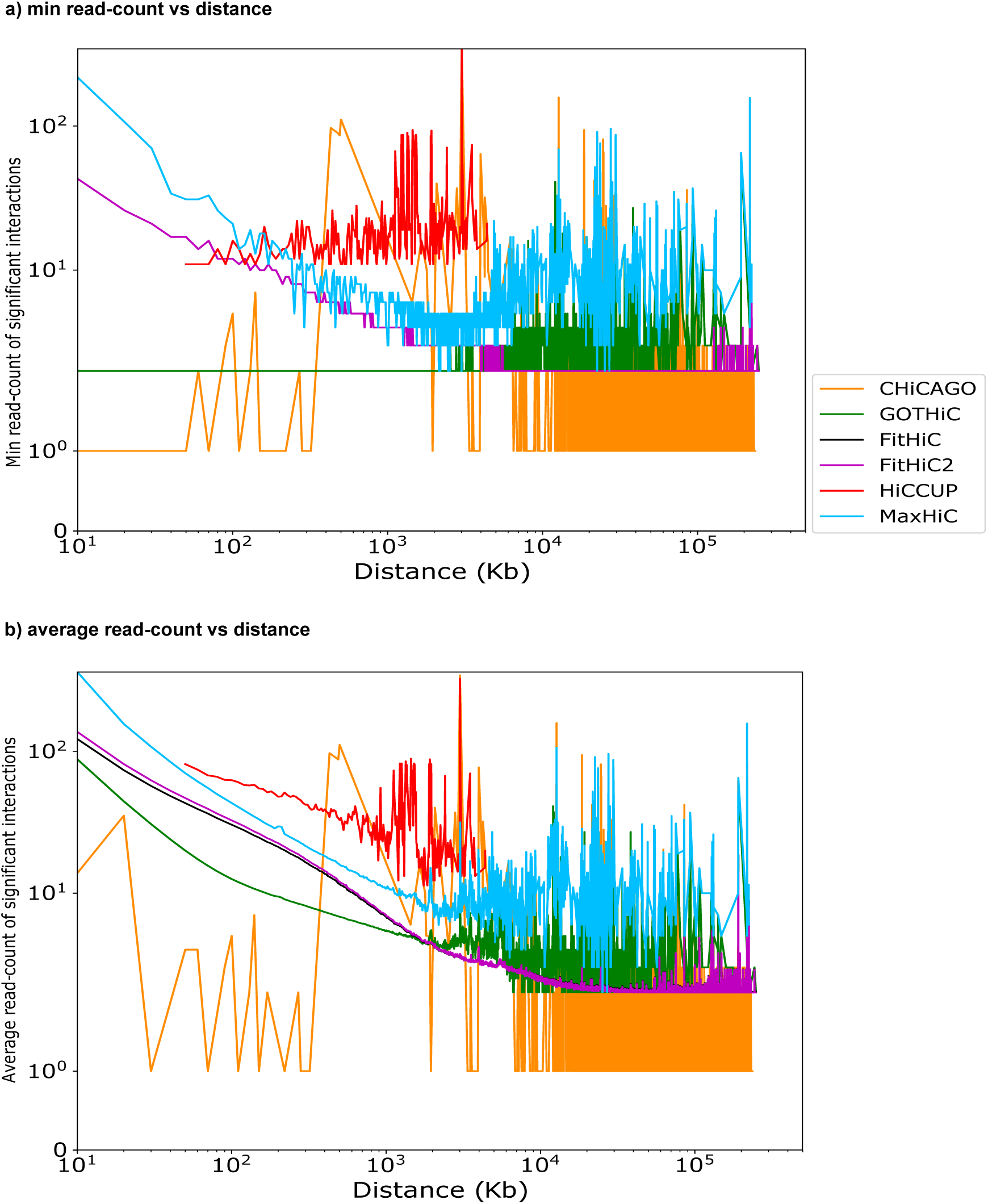
Distance distribution of read-count support of significant interactions identified by the four models in GM12878 and HMEC. **a)** Min read-count of significant interactions in the Rao *et al.* GM12878 data, **b)** as in a but for HMEC, **c)** Average read-count of significant interactions in the Rao *et al.* GM12878 data and **d)** as in c but for the HMEC data. Fragment size 10kb.

**Figure S4.**
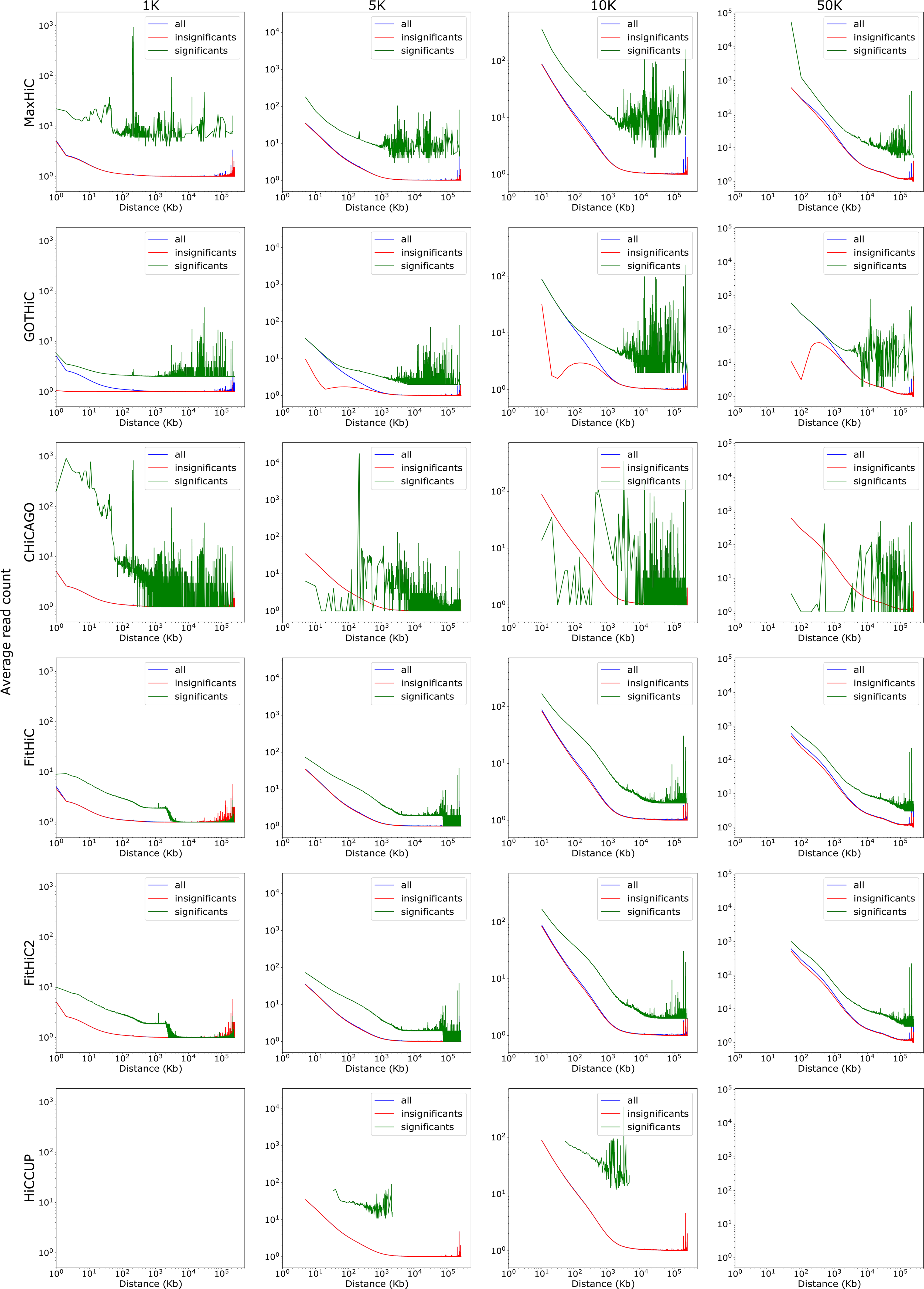
Comparisons of the number of significant interactions and average read-counts of significant interactions identified at fragment sizes 1kb, 5kb, 10kb, 50kb. Plots as in **Figure1.** Note: HiCCUPS did not work on fragment size 1kb and did not identify any significant chromatin loops using fragment size 50kb.

**Figure S5.**
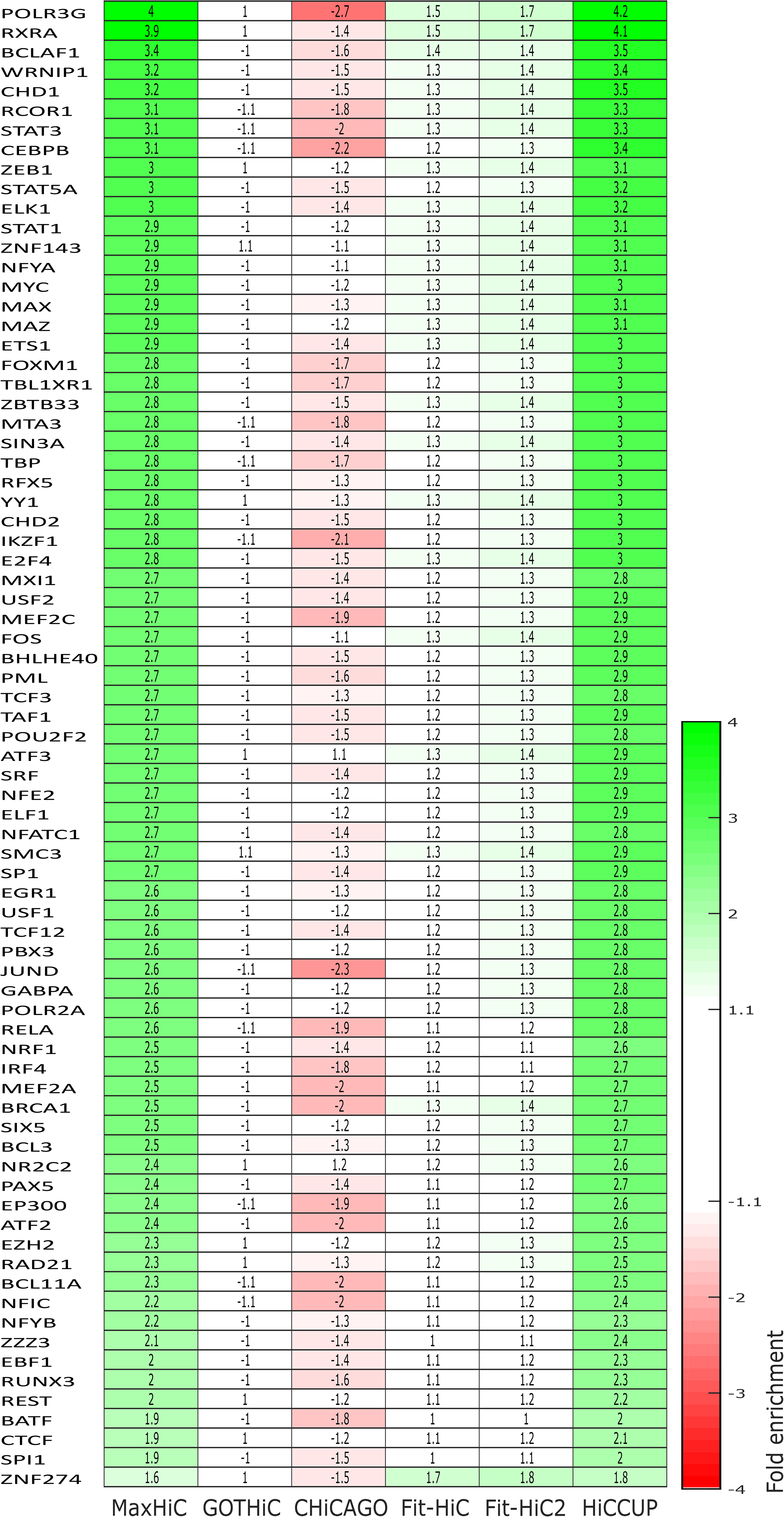
Enrichment of transcription factor binding sites from ENCODE ChIP-seq data on Rao *et al*. GM12878 data. As the plot shows the enrichments in MaxHiC’s interactions are similar to the enrichments in the chromatin loops identified by HiCCUPS (which identified 34 fold fewer interactions).

**Figure S6.**
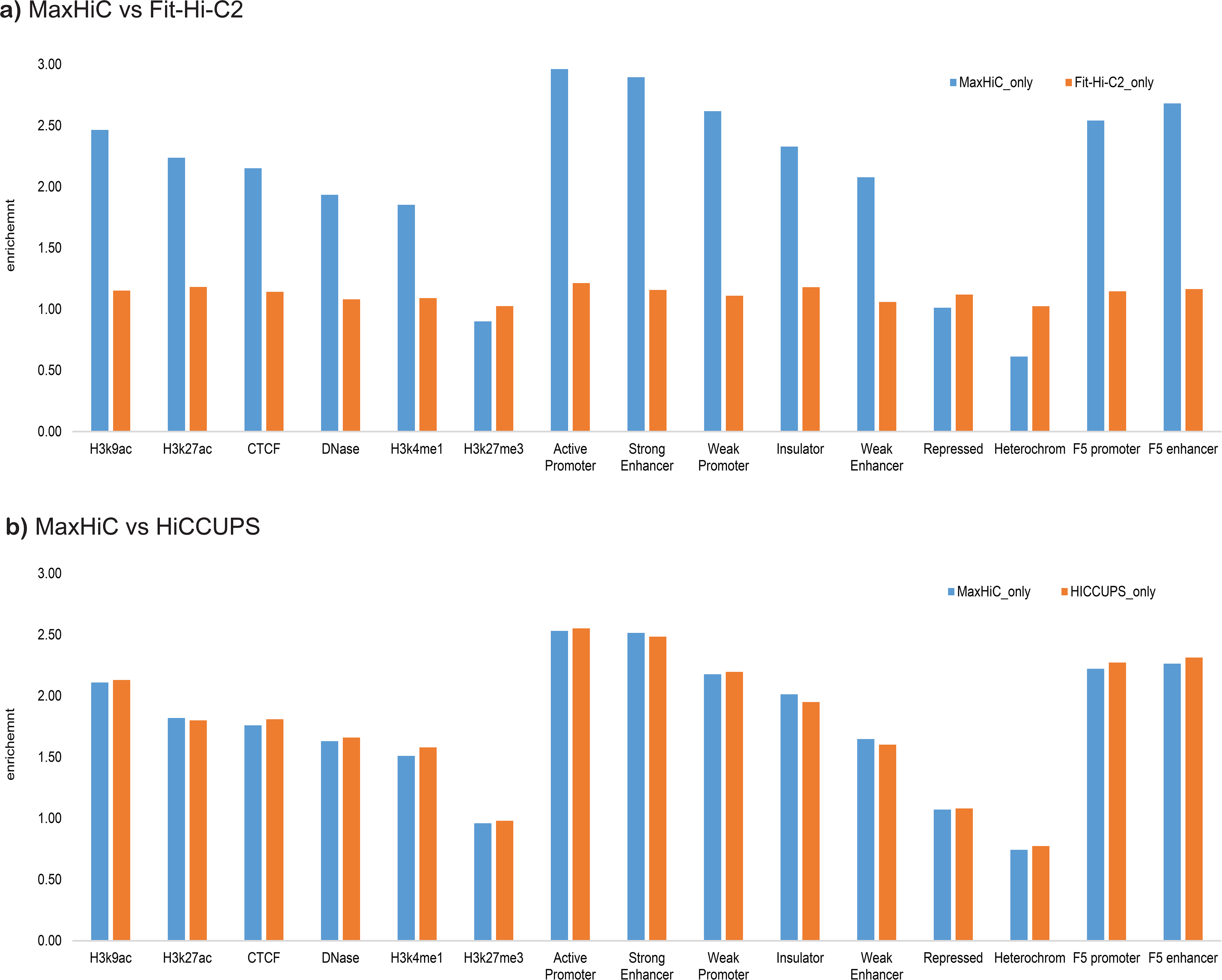
Enrichment of regulatory features in model specific interacting regions on Rao *et al*. GM12878 Hi-C data. a) Interacting regions identified by MaxHiC only versus interacting regions identified by Fit-Hi-C2 only. HICCUPS was excluded from this plot as there was only 1 interacting region identified by HICCUPS and missed by two other tools. The number of MaxHiC-specific interacting regions is 39,629; and the number of Fit-Hi-C2-specific interacting regions is 516,719. b) **Interacting regions identified by MaxHiC only versus interacting regions identified by HiCCUPS.** The number of MaxHiC-specific interacting regions is 219,397; and the number of HICCUPS-specific interacting regions is 61. P-value < 0.001 and fragment size = 10kb.

**Figure S7.**
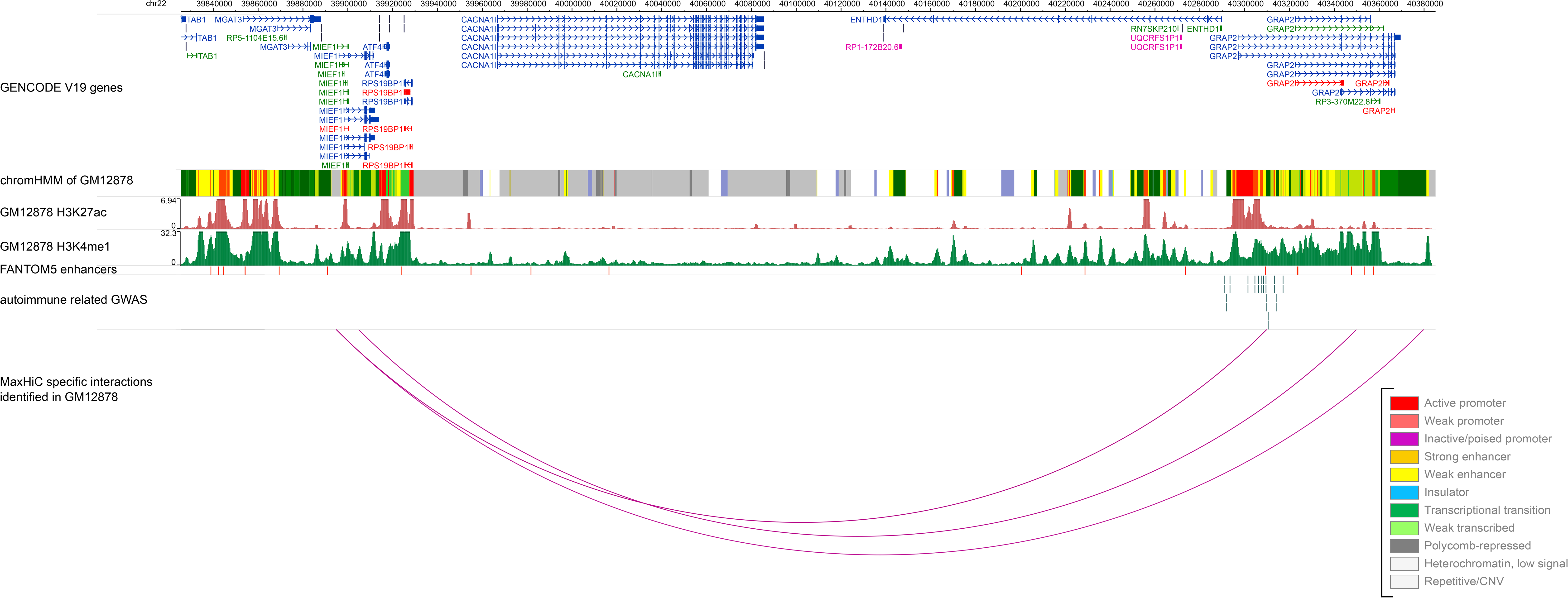
An example of Hi-C interaction identified by MaxHiC as significant, which was not identified as a significant interaction by other tools. Both sides if the interaction have strong signals of regulatory features.

**Figure S8.**
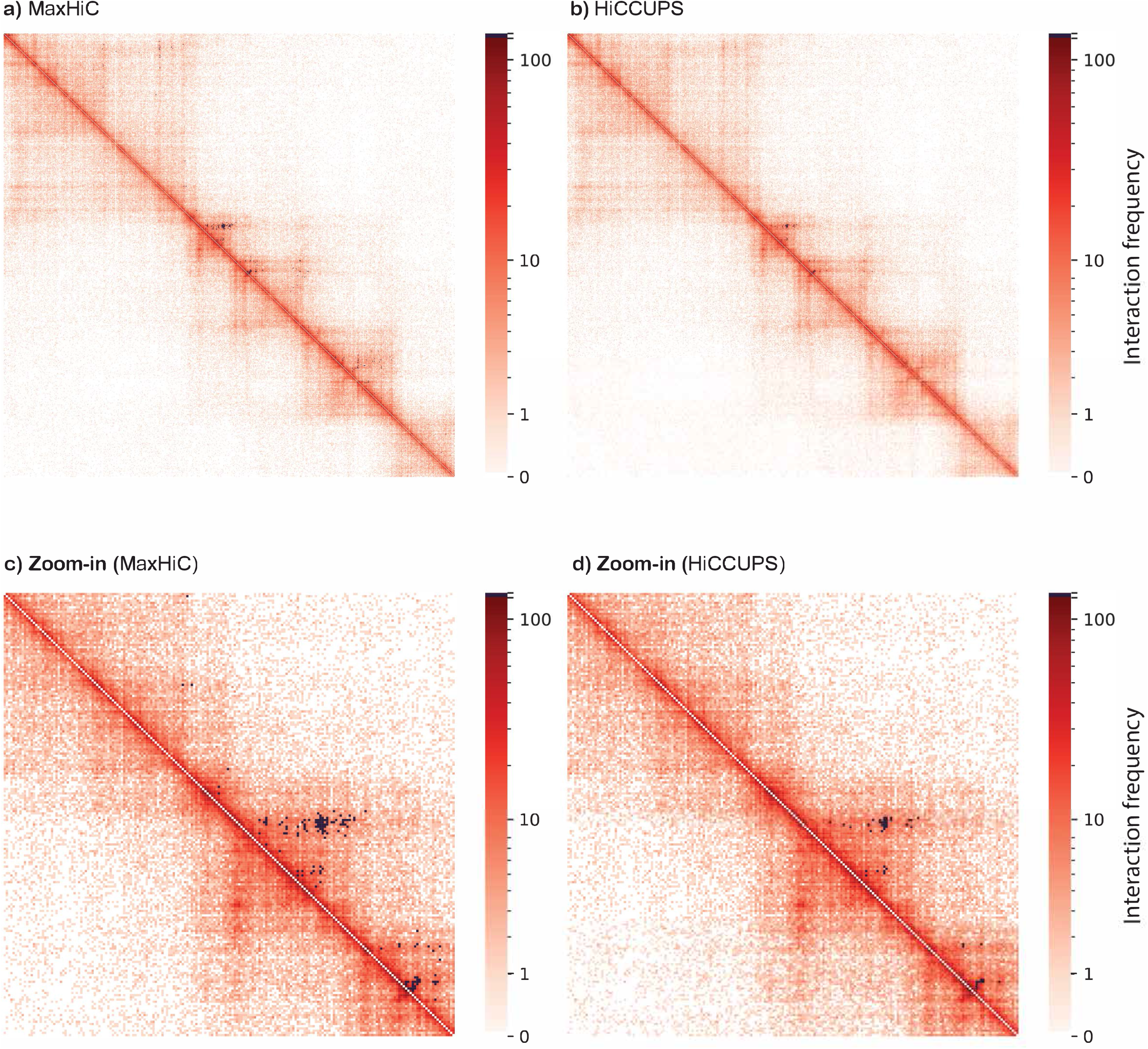
Hi-C interaction map on GM12878 HiC data. Heatmap showing the raw read count of interactions at 5k bins of chr21 (36000kb to 39500kb) (same region **as Supplementary Figure 10 in** the cLoops paper (38) on the GM12878 Hi-C dataset in Rao *et al*. (7). The heatmap is colored based on the log of read count. Significant interactions identified by **a)** MaxHiC and **b)** HICCUPS, are shown in black. A zoomed view of one TAD is shown in **c)** and **d)**.

**Figure S9.**
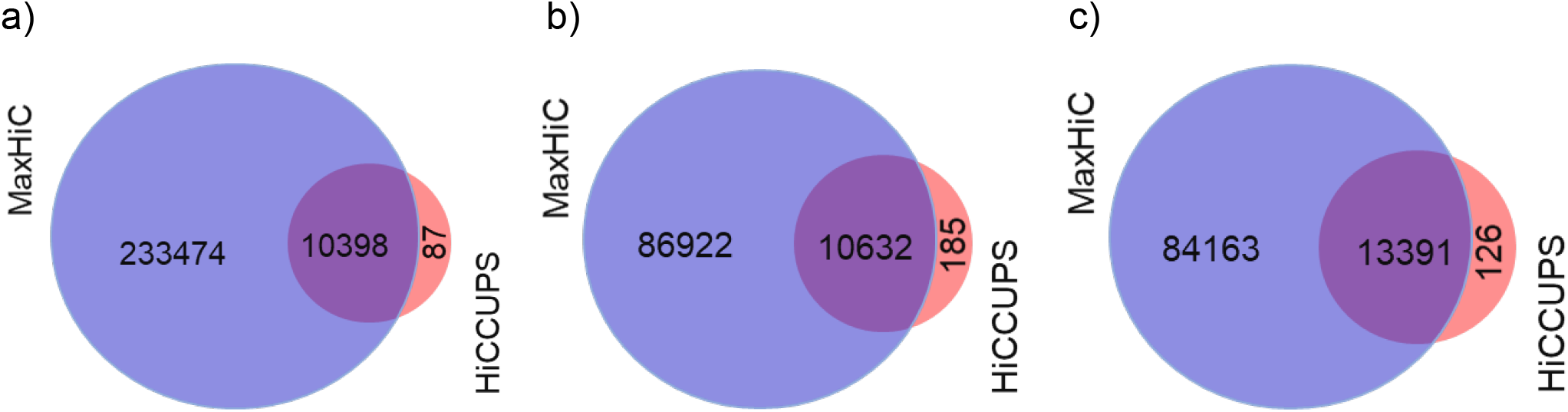
Venn diagrams showing **a)** Overlap between the sets of interactions identified by MaxHiC and HiCCUPS on the GM12878 dataset. **b)** Overlap of 10kb genomic windows involved in significantly interacting regions identified (Note: for HICCUPS only the 10kb centroids are counted). **c)** Similar to b, but showing overlap when the entire anchor sequence is considered for HICCUPS.

**Figure S10.**
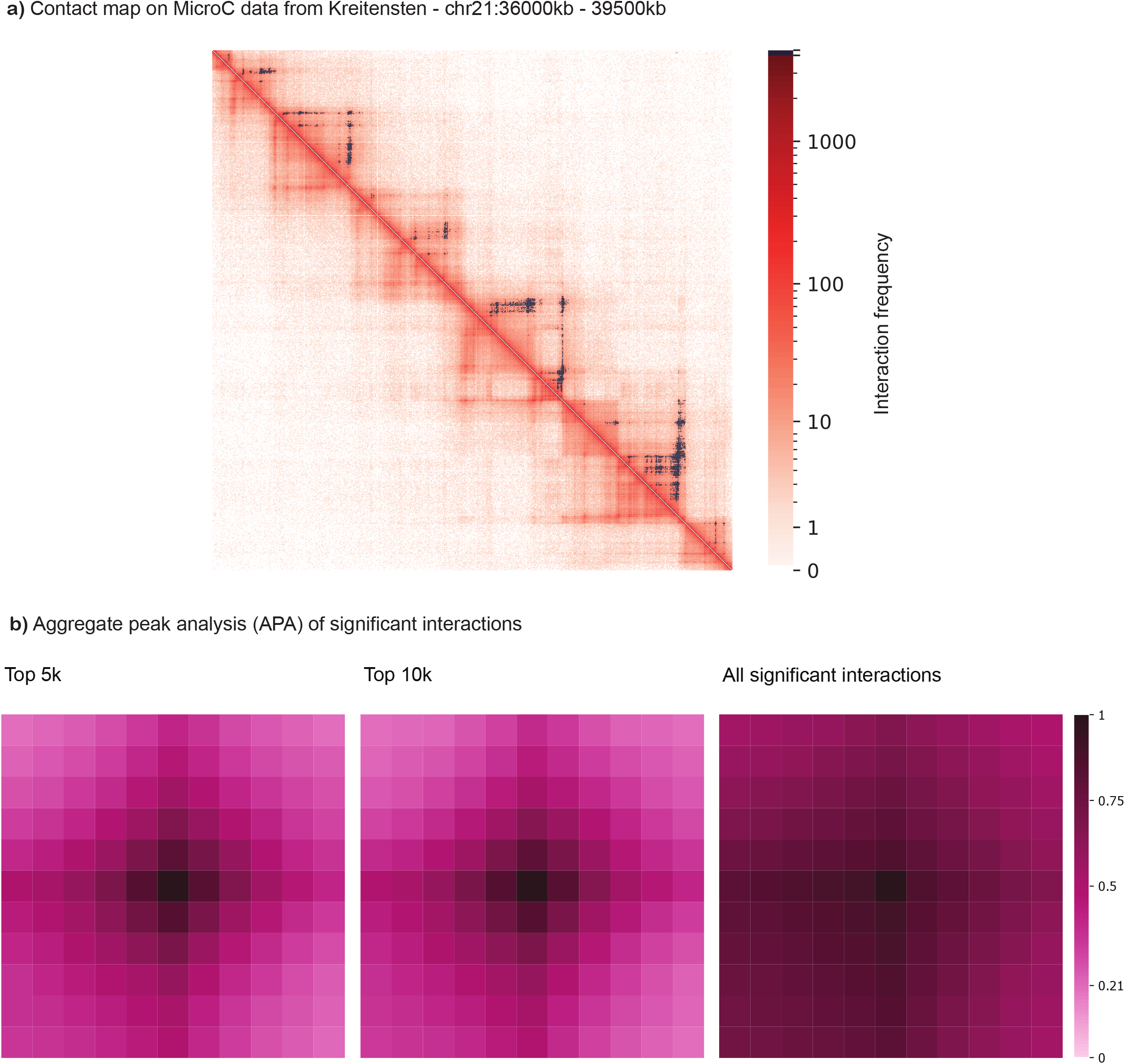
MicroC data analysis. a) Contact map on MicroC data from Kreitensten *et al*. (chr21:36000kb-39500kb). **b) Aggregate peak analysis (APA) of significant interactions called by MaxHiC on MicroC data**. Interactions were called at 5kb resolution using the microC Hi-C dataset from. The analysis shows the aggregate background interaction frequency of 5 bins upstream and downstream of the significantly interacting bins. APA plots are shown for the top 5k, and 10k.

**Figure S11.**
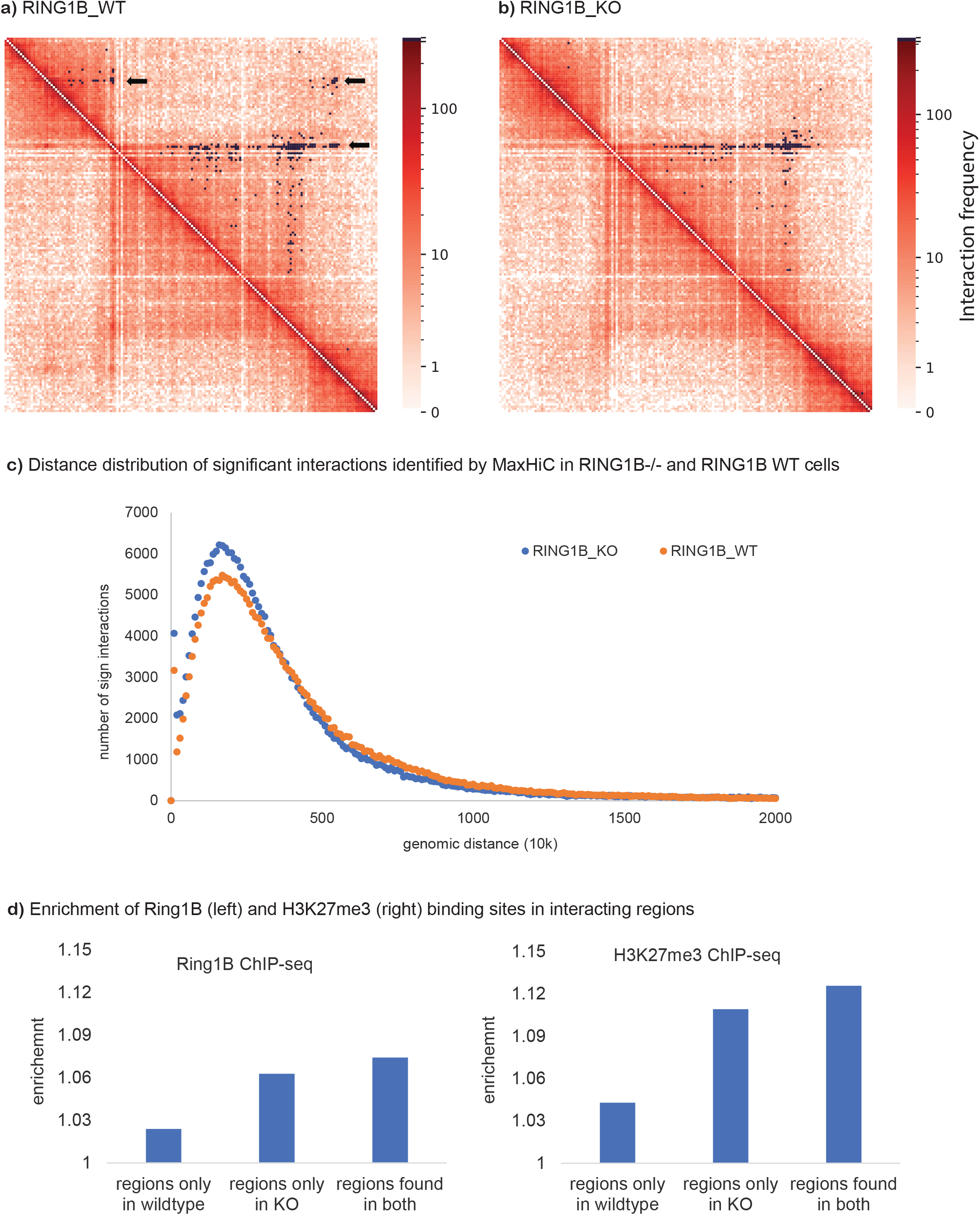
RING1B-depleted Hi-C analysis. a) Hi-C Contact map comparing significant interactions identified in WT and RING1B−/− cells at 10kb fragment size. Black pixels are significant interactions identified by MaxHiC at P-value <0.001. chr5:28.3-30Mb). **b) Distance distribution of significant interactions identified by MaxHiC (Pvalue <0.001, 10kb bins) in RING1B−/−, and RING1B WT cells. c) Enrichment of Ring1b (left) and H3K27me3 (right) binding sites in interacting regions** found i) only in the wild type cells, ii) only observed in the RING1B knockout cells and iii) those stably observed in both.

**Figure S12.**
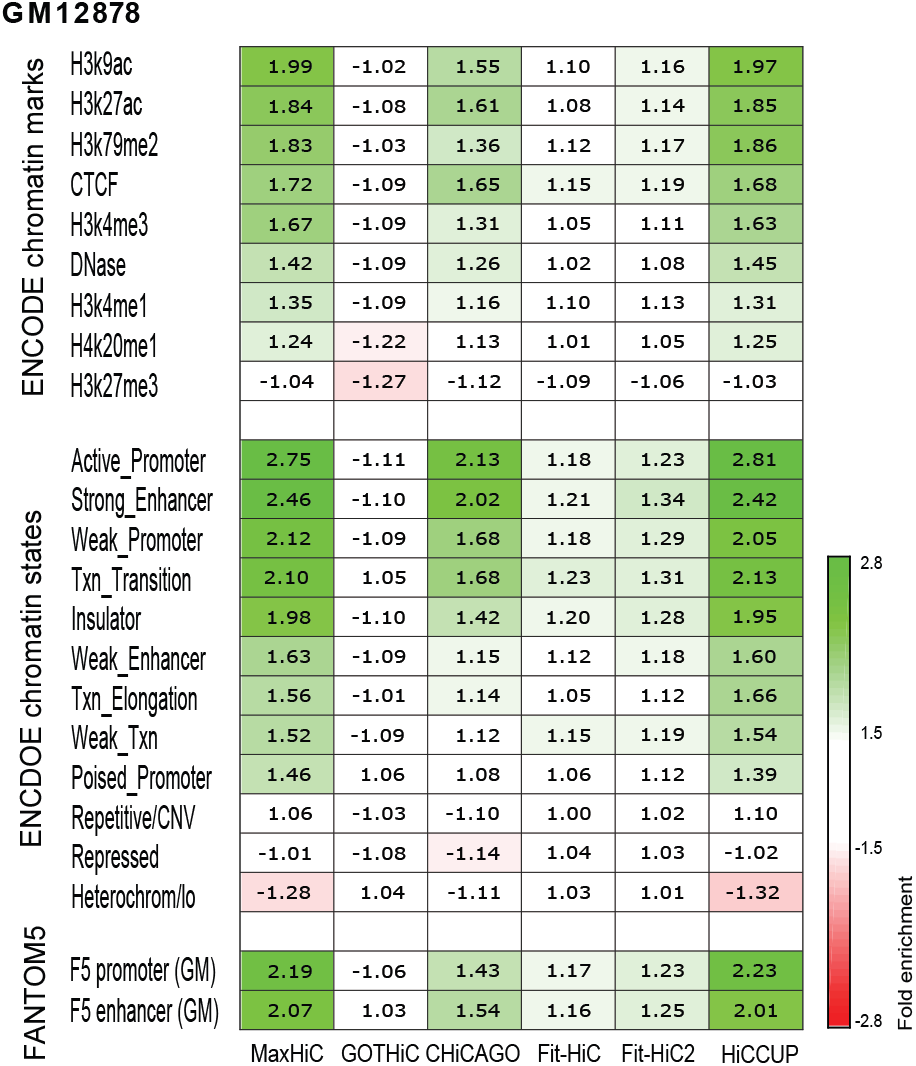
Comparison of the four models in the analysis of capture Hi-C experiment in capture Hi-C sample GM12878. We also developed a separate version of MaxHiC to handle capture Hi-C libraries. In the capture Hi-C library, we are particularly interested to identify significant interactions that at least one end of interaction is baited (refer to capture regions). Enrichment of GM12878 specific histone marks, putative regulatory regions and GWAS signal in regions identified by CHiCAGO, GOTHiC, Fit-Hi-C, Fit-Hi-C2, HiCCUPS and capture version of MaxHiC, on the capture Hi-C library GM12878 from Mifsud *et al.* (13). This showed that significant target-interacting pairs identified by MaxHiC and HiCCUPS have much higher enrichment for regulatory pairs and Autoimmune related GWAS SNPs. Importantly, MaxHiC identified 36 fold more interactions than HiCCUPS but at the same level of enrichments for regulatory features.

## Supplementary tables legends

**Supplementary table S1.** Details of publicly available Hi-C samples used in this study.

**Supplementary Table S2.** The statistical summaries of significant interactions identified by MaxHiC, GOTHiC, CHiCAGO, Fit-Hi-C, Fit-Hi-C2, and HiCCUPS on the Rao HMEC Hi-C library at fragment sizes 1kb, 5kb and 10kb.

**Supplementary Table S3.** Enrichment of transcription factor binding sites in the significant interactions identified by MaxHiC, GOTHiC, CHiCAGO, Fit-Hi-C, Fit-Hi-C2, and HiCCUPS on the Rao GM Hi-C library at fragment sizes 10kb.

**Supplementary Table S4.** Details of autoimmune and breast cancer traits used in this study for GWAS analysis.

**Supplementary Table S5.** The statistical summaries of significant interactions identified by MaxHiC, GOTHiC, CHiCAGO, Fit-Hi-C, Fit-Hi-C2, and HiCCUPS on the Mifsud GM12878 capture Hi-C library at fragments 5kb and 10kb. As the table shows, the average read count of significant interactions identified by MaxHiC are much higher than those identified by CHiCAGO, GOTHiC, Fit-Hi-C, Fit-HiC2 and HiCCUPS.

